# Conditional Deletion of All Neurexins Defines Diversity of Essential Synaptic Organizer Functions for Neurexins

**DOI:** 10.64898/2026.07.15.738748

**Authors:** Man Jiang, Yang Li, Margarita Artyukhova, Li Zhang, Shu Luo, Weiran Zhan, Bo Zhang, Ozgun Gokce, Thomas C. Südhof

## Abstract

Presynaptic neurexins are key regulators of synapse properties that arguably represent the best-studied synaptic adhesion molecules. Despite thousands of papers, however, no direct comparison of overall neurexin functions in different types of synapses is available. A decade ago, we provided such an analysis but we recently retracted this paper because four images contained microduplications that, although without discernible impact on the paper’s conclusions, could not be corrected. As a result, the scientific community lost access to primary data that established the fundamental principle that neurexins perform profoundly different functions in different types of synapses. In the present study, we have therefore reanalyzed the original raw data and expanded their conclusions with new experiments to document, in a single study, the basic contributions of neurexins to different synapses. We used triple conditional knockout mice targeting all neurexins except for Neurexin-1γ and applied neuron-specific manipulations combined with slice electrophysiology, two-photon Ca^2+^-imaging, and immunohistochemistry. With this approach, we focused on excitatory climbing-fiber synapses in the cerebellum and on inhibitory synapses formed by parvalbumin- or somatostatin-positive neurons in the cerebellum, hippocampus, and medial prefrontal cortex. Our results show that pan-neurexin deletions produce dramatically different phenotypes in synapses, ranging from modest to massive impairments in synapse assembly (climbing-fiber and parvalbumin-positive synapses) to severe but selective decreases in action potential-induced presynaptic Ca^2+^-transients (somatostatin-positive synapses). Thus, neurexins perform powerful but distinct context-dependent roles in different synapses that shape the brain’s circuits.

**Significance Statement:** Neurexins are abundant presynaptic adhesion molecules expressed from three genes in thousands of splice variants. Although neurexins are well studied, no analysis that compares neurexin deletions in multiple types of synapses is currently available. Here, we provide such an analysis by examining triple conditional knockout mice that delete all neurexins except for Neurexin-1γ. Using neuron-specific manipulations combined with slice electrophysiology, two-photon Ca^2+^ imaging and immunohistochemistry, we show that pan-neurexin deletions produce distinct phenotypes in different synapses, ranging from impairments in synapse assembly (climbing-fiber and parvalbumin-positive cortical synapses) to severe selective decreases in presynaptic action potential-induced Ca^2+^-transients (somatostatin-positive cortical synapses). Our data reveal context-dependent distinct functions of neurexins in different synapses whose properties shape the input-output relations of neural circuits.

## Introduction

Neurexins are presynaptic cell-adhesion molecules that were identified as α-latrotoxin receptors (1–4). In vertebrates, three neurexin genes (*Nrxn1*, *Nrxn2*, and *Nrxn3* in mice) encode longer α- and shorter β-neurexins from independent promoters (3, 5). In addition, the *Nrxn1* gene encodes a third, even shorter γ-neurexin also using an internal promoter (6). Neurexins are evolutionarily conserved. A single α-neurexin is expressed in Drosophila (5) and single α- and γ-neurexins are present in C. elegans (7). Moreover, broadly defined neurexin-related genes, such as CNTNPs, are present throughout the animal kingdom, including in cnidaria that contain only a rudimentary nervous system (8, 9).

α-Neurexins are composed of a large extracellular sequence comprising six LNS-domains (for laminin/neurexin/sex-specific globulin-domain) with three interspersed EGF-like domains, followed by an O-linked carbohydrate attachment sequence, a single heparan-sulfate attachment site (10), a cysteine-loop domain (6), a transmembrane region, and a short cytoplasmic sequence (1) (Figure S1A). β-Neurexins include a short β-neurexin-specific N-terminal sequence that then splices into the α-neurexin sequence N-terminal to the last, sixth LNS-domain; from that point on, α- and β-neurexins are identical (3). Neurexin-1γ, finally, also contains a unique short N-terminal sequence that then splices into the α- and β-neurexin sequences C-terminal to the last, sixth LNS-domain, again with identical C-terminal sequences (6) (Figure S1A).

Individual neurexins are essential for multiple synaptic functions in C. elegans, Drosophila, mouse, and human neurons, but no uniform overall role for neurexins has emerged (7, 11–41). Genetic mutations of neurexin genes, analyzed in various animals and different synapses, uncovered diverse phenotypes that affect multiple aspects of synaptic function. In mice, constitutive triple KO of all α-neurexins is lethal (11). At birth, triple α-neurexin KO brains exhibit a modest decrease in inhibitory but not excitatory synapse density and display dramatic impairments in excitatory and inhibitory synaptic transmission due, at least in part, to a loss in presynaptic Ca^2+^-influx but do not feature impairments in axon or dendrite outgrowth (11, 32, 33, 42, 43). Moreover, triple α-neurexin KO mice exhibit an impairment in neuropeptide signaling (44).

Triple β-neurexin KO mice, moreover, also display a survival phenotype, albeit a milder one than that of triple α-neurexin KO mice, consistent with the relatively lower expression of β-neurexins than α-neurexins (21, 45). Furthermore, triple β-neurexin KO mice exhibit a synaptic transmission impairment that is due, at least in part, to a disinhibition of tonic endocannabinoid signaling (21) and a robust decrease in the number of presynaptic dense-core vesicles, again suggesting functions in neuropeptide signaling (46).

Triple α/β-neurexin KO mice that target both α- and β-neurexins, finally, were reported by us in 2017 in a paper that we retracted because four images contained inexplicable microduplications that did not materially affect the conclusions of the paper (47, 48). The images could not be corrected because the original samples were no longer retrievable after a decade, although other raw data were available. The Chen et al. (2017) paper (47) was foundational because it demonstrated that the triple α/β-neurexin deletions (called pan-neurexin deletions) produced distinct phenotypes in different synapses without abolishing synapse formation, suggesting that neurexins are not essential for synapse formation as such but shape synapses in a context-specific manner. This conclusion was expanded in subsequent studies showing among others that, in excitatory Calyx of Held synapses and in synapses formed by MNTB neurons on the superior olive, the conditional triple α/β-neurexin KO produced a selective decrease in action potential-triggered Ca^2+^ influx at the active zone (49, 50). Additional studies reported that the conditional triple α/β-neurexin KO induces a general disorganization of presynaptic active zones that includes impairments in presynaptic receptors (51), decreases Ca^2+^-triggered exocytosis in dopaminergic neurons (52), and may cause neuronal cell death in olfactory sensory neurons, cerebellar granule cells, and serotonergic neurons of the brain stem (53–55).

Thus, present analyses of the global functions of neurexins suggest important and robust synaptic and possibly non-synaptic roles in different types of neurons. However, no direct comparison of α/β-neurexin KO-induced phenotypes in different types of neurons and synapses is currently available after retraction of our original study (47). Moreover, in many studies the phenotypes were examined primarily morphologically without functional analyses of synaptic transmission, rendering the nature of observed changes unclear. To address these knowledge gaps at least in part, we here aimed to specifically clarify whether neurexins perform canonical, broad functions in all synapses or synapse-specific functions that depend on the pre- and postsynaptic neuron types. For this purpose, we reanalyzed the raw data of our retracted Chen et al. (2017) paper (47) and performed additional experiments to complement the original available data, with all raw data made publicly available for independent assessments (website link: https://purl.stanford.edu/xp149zq3345). As a result, the present study reports an electrophysiological and morphological analysis of the effect of the triple α/β-neurexin conditional deletion on multiple well-defined synapses, namely excitatory climbing-fiber synapses in the cerebellum and inhibitory synapses formed by parvalbumin-positive (Pv^+^) or somatostatin-positive (SST^+^) interneurons on hippocampal CA1 neurons and on layer 5 pyramidal neurons in the medial prefrontal cortex (mPFC). Our data reveal that the presynaptic triple neurexin deletions produce two distinct principal types of phenotypes in the examined synapses, namely a loss of synapses or a decrease in voltage-gated presynaptic Ca^2+^ influx, both of which result in a decrease in synaptic connectivity. Thus, our results indicate that neurexins perform functionally diverse fundamental roles in different types of synapses.

## Results

### Design and objective

The primary goal of the present study is to compare the effects of deleting all neurexins except for Nrxn1γ across different, well-defined types of synapses in multiple brain regions. For this purpose, we quantitatively reanalyzed the existing raw data of our previous paper (47) that we retracted because of discrete image abnormalities (48) and performed additional experiments to complement the previously obtained data (see Experimental Design, Figure 1B), with public posting of all raw data for access to the community (SDR weblink: https://purl.stanford.edu/xp149zq3345). This effort was made to ensure that the results from our original foundational study (47) would not be lost for the scientific community and that the conclusions of this paper could be validated by new independent analyses.

**Figure 1.**
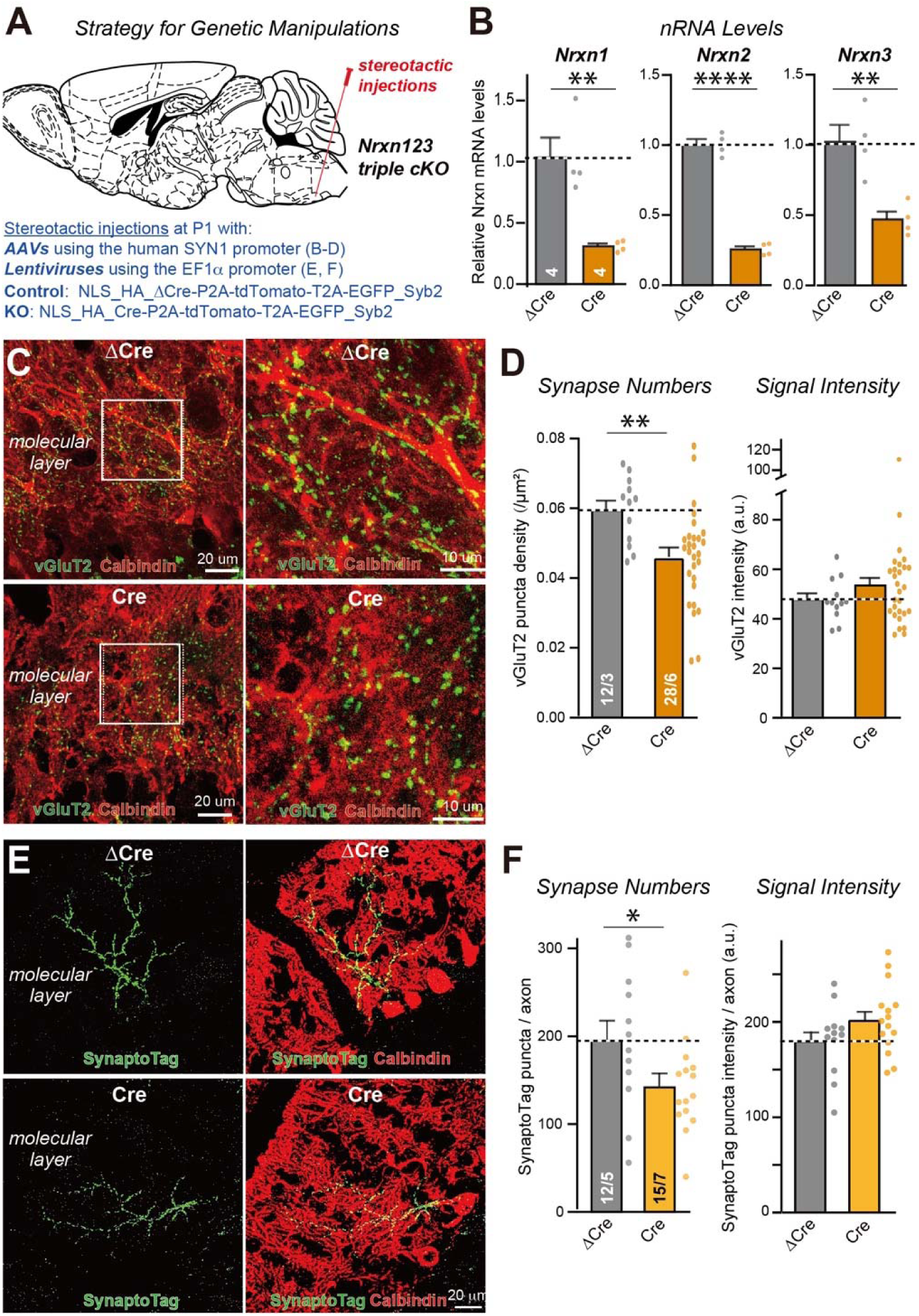
AAV-mediated global and lentiviral sparse pan-neurexin deletions in the inferior olive significantly lower the climbing-fiber synapse density. **A**, Schematic of the stereotactic ablation of neurexins in the inferior olive at P1 by AAV-mediated or lentiviral expression of mutant inactive (ΔCre, control) or wild-type active Cre-recombinase. Analyses were performed at P28. **B**, qRT-PCR measurements of neurexin mRNA levels in the inferior olive after AAV-mediated delivery of ΔCre or Cre. **C**, Representative images of the cerebellar cortex of AAV-injected mice visualizing Purkinje cells by calbindin (red) and climbing-fiber synapses by vGluT2 immunofluorescence (green) (left, low-magnification images; right, high-magnification images from the boxed areas). **D**, Quantification of climbing-fiber synapse numbers (left) and vGluT2 staining intensity (right). **E**, Representative images of the cerebellar cortex from injected mice visualizing Purkinje cells by calbindin immunofluorescence (red) and synaptic specializations formed by infected inferior olive neurons by SynaptoTag labeling (green, Synaptobrevin 2-EGFP). **F**, Quantification of the climbing fiber synapse numbers per climbing fiber (left) and SynaptoTag signal intensity (right). Summary graphs show means ± SEMs; numbers in graphs indicate the number of mice (B), the number of sections/mice (D), or the number of climbing fibers/mice (F) (* = p<0.05; ** = p<0.01; **** = p<0.0001 by Student’s t-test).

### Presynaptic neurexin deletion from climbing-fiber synapses modestly impairs synapse assembly

To analyze the effect of deleting neurexins from climbing-fiber synapses, we pursued three complementary approaches: Injection of AAVs expressing mutant (ΔCre, control) or wild-type Cre-recombinase or of lentiviruses co-expressing Synaptobrevin2-EGFP (SynaptoTag) tracers (56) with mutant (ΔCre, control) or wild-type Cre-recombinase into the inferior olive of newborn *Nrxn123* triple cKO mice (Figure 1A, S1B1-2), and crossings of *Nrxn123* triple cKO mice with Crh-Cre driver mice that selectively express Cre-recombinase in inferior olive neurons (57) (Figure 2, S1B3).

**Figure 2.**
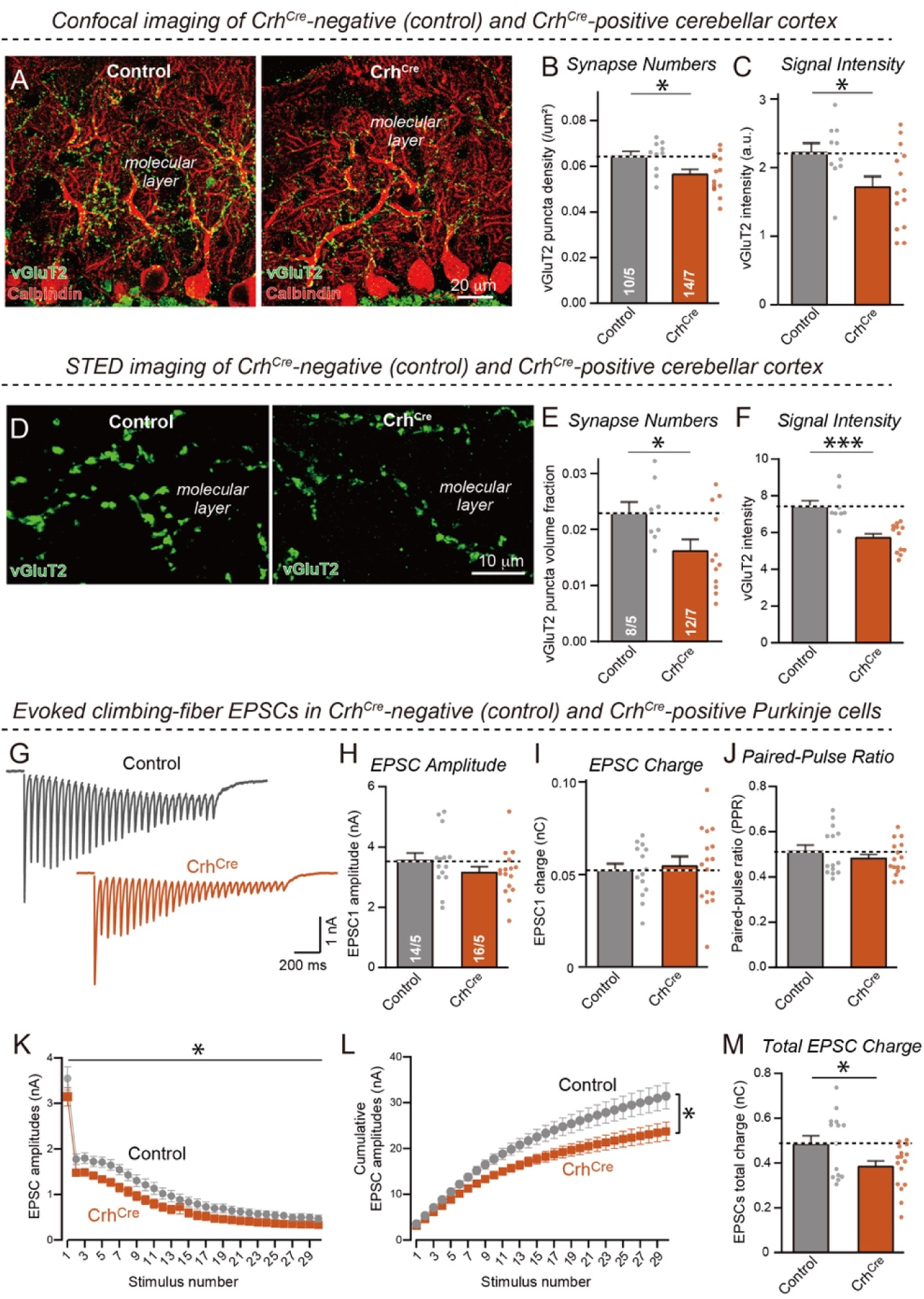
Crh-Cre-mediated global pan-neurexin deletions in the inferior olive significantly decrease climbing-fiber synapse numbers. **A**, Representative confocal images of cerebellar cortex sections from triple *Nrxn123* cKO mice without (control) or with a Crh-Cre allele mediating Cre-expression in the inferior olive. Sections were stained at P28 for vGluT2 (green) and calbindin (red). **B** & **C**, quantifications of the density (B) and staining intensity (C) of vGluT2-positive climbing-fiber synapses) as a function of the Crh-Cre allele. **D** -**F**, Analysis by STED super-resolution microscopy of the effect of the Crh-Cre mediated pan-neurexin deletion in the inferior olive on vGluT2-positive climbing-fiber synapses (**D**, representative STED images of cerebellar cortex sections stained for vGluT2 (green); **E** & **F**, quantifications of the vGluT2 puncta volume fraction as a proxy for synapse density (E) and of the vGluT2-staining intensity of climbing-fiber synapses (F)). **G**, Representative traces of climbing-fiber EPSCs during a 20-Hz stimulus train recorded from Purkinje cells in acute cerebellar slices of triple *Nrxn123* cKO mice lacking or harboring a Crh-Cre allele (performed at P33-37 in picrotoxin and a low dose of CNQX. **H** - **J**, Quantifications of EPSC1 amplitude (H), EPSC1 charge transfer (I), and paired-pulse ratio (PPR) at 50 ms (J). **K** & **L**, EPSC amplitudes (K) and cumulative EPSC amplitudes (L) as a function of stimulus number. **M**, Quantifications of the total EPSC charge transfer during a 20-Hz stimulus train. All numerical data are means ± SEMs (n’s (sections/mice or neurons/mice) are indicated in graphs); statistical evaluations were performed by Student’s t-test or two-way ANOVA with post-hoc corrections (K, L) with * = p<0.05; *** = p<0.001).

In the first approach, we administered AAVs by stereotactic delivery into the inferior olive of newborn mice and analyzed the mice at P28 (Figure 1A). The AAVs broadly infected olivary neurons, enabling us to measure the efficacy of the Cre-mediated deletion by quantitative RT-PCR. AAV-mediated Cre-recombination in neurons decreased *Nrxn1* and *Nrxn2* mRNAs by ∼75% and *Nrxn3* mRNAs by ∼55% (Figure 1B). These measurements are likely underestimates since neurexins are also expressed by astrocytes and since mutant neurexin mRNAs may not be fully degraded by nonsense-mediated decay. We then analyzed climbing-fiber synapses in the cerebellar cortex as a function of the pan-neurexin deletion. Immunocytochemistry for vGluT2, a specific marker for these synapses (58), revealed a significant but limited decrease in climbing-fiber synapse density (∼25%) without a change in the vGluT2 staining intensity per punctum (Figure 1C, D).

In the second approach in which we infected a subset of inferior olive neurons of newborn mice with lentiviruses co-expressing SynaptoTag (56) with ΔCre or Cre, we again analyzed the mice at P28 (Figure 1A). The lentivirus produced sparse co-expression of ΔCre or Cre with SynaptoTag. Climbing fibers from infected neurons were identified in the cerebellar cortex by virtue of their SynaptoTag fluorescence (Figure 1E, F; S2A, B). Deletion of neurexins again caused a significant decrease (∼30%) in climbing-fiber synapse numbers on proximal dendrites of cerebellar Purkinje cells without changing the SynaptoTag signal intensity (Figure 1F).

In the third approach, we independently assessed the effect of pan-neurexin deletions in the inferior olive using a non-viral method that analyzed littermate pan-neurexin cKO mice containing or lacking a Crh-Cre driver allele, which expresses Cre-recombinase in inferior olive neurons (57). Analysis of the effect of the pan-neurexin deletions induced by Crh-Cre in inferior olive neurons on climbing-fiber synapses using immunohistochemistry for the climbing-fiber-specific marker vGluT2 (58) confirmed the decrease (∼30%) in synaptic puncta numbers observed with the viral pan-neurexin deletions (Figure 2A-C; S2C). This conclusion was further validated by quantitative STED super-resolution microscopy, which reproduced the ∼30% loss of synapses caused by the *Nrxn123* triple deletion. With both types of imaging, the remaining synapses exhibited a decrease in the vGluT2 staining intensity (∼20-30%) (Figure 2D-F), which we had not observed for the AAV-mediated *Nrxn123* triple deletion.

The expression of Cre-recombinase in all inferior olive neurons in Crh-Cre driver mice enabled additional analyses of the effects of the neurexin deletions by electrophysiology (Figure 2G-M; S3). Consistent with the morphological results, the *Nrxn123* triple deletion produced a modest decrease in synaptic connectivity, with a non-significant trend towards a decline in the amplitude of single evoked EPSCs that became significant upon analyses of trains of EPSCs induced at 20 Hz (Figure 2G-M; Figure S3).

Viewed together, these data uncover a reproducible but limited impairment of climbing-fiber synapse assembly upon deletion of neurexins from inferior olive neurons. This phenotype is similar to, but less severe than, the previously observed phenotype (47), possibly because the previous experiments suffered from viral toxicity or because the present experiments may not have induced as complete a neurexin deletion as the previous experiments, as also indicated by the qRT-PCR measurements of the neurexin mRNAs after the AAV-mediated deletion (Figure 1B).

### The pan-neurexin deletion impairs the organization of inhibitory synapses formed by hippocampal Pv^+^ interneurons

To analyze the effect of the pan-neurexin deletion on inhibitory synapses, we crossed *Nrxn123* triple cKO mice with Pv-Cre mice that express Cre-recombinase under the parvalbumin promoter (Figure S1B4). We then cut hippocampal sections from adult littermate *Nrxn123* cKO/Pv-Cre and *Nrxn123* cKO mice and analyzed the sections by immunohistochemistry for vGAT as a marker of inhibitory synapses and for parvalbumin as a marker of the targeted inhibitory interneurons (Figure S4A). Since confocal imaging could not resolve individual synaptic puncta in the stained sections, we quantified the overall synapse density as the area fraction occupied by vGAT fluorescence (Figure S4B). The pan-neurexin deletion in Pv^+^ neurons produced a ∼50% decrease of the vGAT signal in the S. oriens and S. radiatum of the CA1 region and a lesser ∼20% decrease in the S. pyramidale (Figure S4B). Given that synapse quantifications by immunohistochemistry depend critically on the intensity and object-sizing thresholds chosen to differentiate signal from background noise, the decrease in synaptic signal observed here is consistent with a decrease in synapse numbers or synapse size.

We next analyzed inhibitory synapses in the hippocampal CA1 region by electrophysiology of adult littermate *Nrxn123* cKO/Pv-Cre and *Nrxn123* cKO mice (Figure S4C-N) but detected no significant phenotype using measurements of spontaneous IPSCs (sIPSCs) (Figure S4C-G) or of input-output curves and paired-pulse ratios of evoked IPSCs (Figure S4H-N). The observed lack of an electrophysiological phenotype here, in view of the immunocytochemically determined decrease in synapse assembly, could be due to several factors, since these experiments measure inhibitory synaptic transmission of all inputs, not just Pv^+^ synapses. Inhibitory synapses on pyramidal neurons from other, non-Pv^+^ interneurons might dominate or a subset of Pv^+^ neurons might form synapses on other inhibitory interneurons (59) that in turn innervate pyramidal neurons, causing a disinhibition of inhibitory responses when Pv^+^ neuron synapses are impaired. Thus, to obtain a clearer analysis, we performed paired recordings in the mPFC as an additional brain region that is relevant for insight into neuropsychiatric disorders.

### The pan-neurexin deletion decreases assembly of inhibitory Pv^+^ interneuron synapses on pyramidal neurons in the mPFC

Our observations in the hippocampus suggested that pan-neurexin deletion impairs synapse assembly, but provided unclear results because overall inhibitory synapses, instead of specifically Pv^+^ synapses, were analyzed. To gain specific insight into Pv^+^ inhibitory synapses and investigate another brain region, we examined the effect of the pan-neurexin deletion on Pv^+^ synapses of the mPFC (Figure 3). Immunohistochemistry for vGAT and parvalbumin again uncovered a significant decrease in inhibitory synaptic signal (∼50%), measured as the vGAT-positive area fraction (Figure 3A, B). Moreover, recordings of miniature IPSCs (mIPSCs) from layer 5 pyramidal neurons of the mPFC revealed a ∼25% decrease in mIPSC frequency without a significant decrease in mIPSC amplitude (Figure 3C-E; Figure S5), consistent with the decrease in synapse assembly.

**Figure 3.**
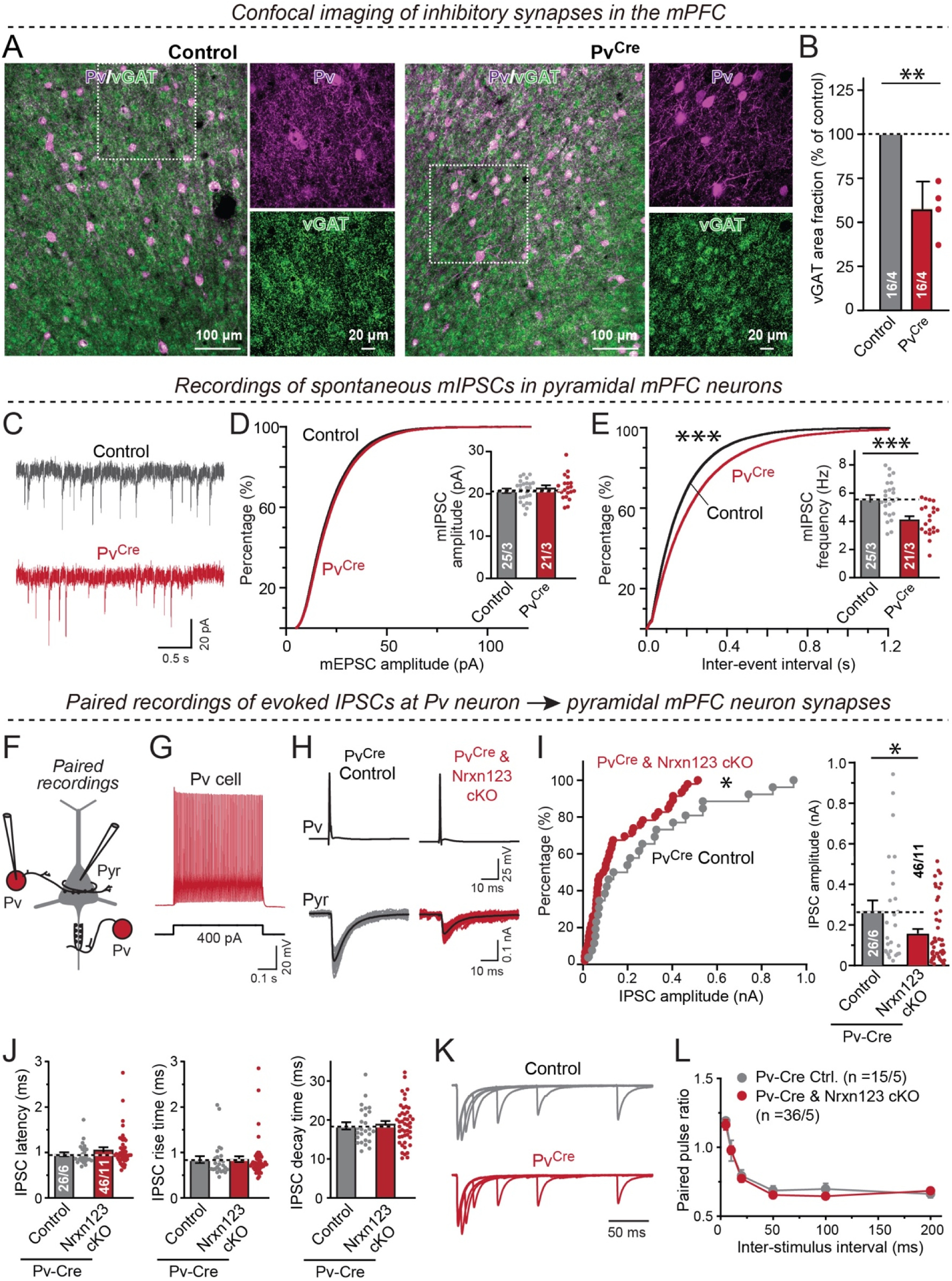
Pan-neurexin deletions in Pv^+^ neurons decrease inhibitory synapse numbers and synaptic transmission in the medial prefrontal cortex (mPFC) **(A)** Representative images of mPFC sections from littermate adult triple *Nrxn123* cKO mice harboring or lacking a Pv-Cre allele. Sections were stained for vGAT (green) and parvalbumin (magenta) (left, overviews; right, zoomed in confocal single-channel images from the boxed areas on the left). **(B)** Quantifications of the area fraction occupied by inhibitory synapses identified by vGAT staining. **(C)** Representative mIPSC traces recorded from mPFC layer 5 pyramidal neurons in acute slices from littermate control *Nrxn123* cKO and *Nrxn123* cKO/ Pv-Cre cells. **(D** & **E)** Cumulative distribution plots of mIPSC amplitudes (left) and inter-event intervals (right) monitored in pyramidal neurons of *Nrxn123* cKO and *Nrxn123* cKO/Pv-Cre mice (insets = summary graphs of average mIPSC amplitude (left) and frequency (right)). (F) Diagram of paired recording configuration (Pv, Pv^+^ interneurons; Pyr, pyramidal neuron). Visualization of Pv^+^ neurons was enabled by viral injections of AAVs encoding double-floxed eYFP into Pv-Cre mice. (G) Representative traces of fast-spiking action potential firing in Pv^+^ interneurons. (H) Representative traces of evoked unitary IPSCs in Pv^+^ interneuron/ pyramidal neuron pairs (gray traces, overlaid individual responses from different trials; black/blue traces, averaged responses). (I) Left, cumulative probability plot of unitary mean IPSC amplitudes (inset = summary graph of IPSC amplitudes) showing that Pv^+^ interneurons from *Nrxn123* cKO/Pv-Cre mice exhibit a decrease in synaptic connectivity. (J) Summary plots of the latency (left), rise time (center), and decay time (right) of unitary IPSCs in Pv-interneuron pyramidal neuron synapses. (K) Representative traces showing PPRs of unitary IPSCs in Pv-interneuron pyramidal neuron synapses at different inter-stimulus intervals. (L) Summary graph of PPRs as a function of the inter-stimulus interval. All numerical data are means ± SEMs (n’s = sections or slices/mice are indicated in the bar graphs); statistical evaluations were performed by Student’s t-test (B-J), Kolmogorov-Smirnov test (cumulative distribution) (D, E, I) or two-way ANOVA followed by Bonferroni *post-hoc* test (L) with * = p<0.05; ** = p<0.01; *** = p<0.001.

Next, we embarked on a mechanistic analysis of the effect of the pan-neurexin deletion on synapses formed by Pv^+^ interneurons onto pyramidal neurons using paired recordings in acute slices (Figure 3F). This approach avoids the inherent limitation of overall synaptic response recordings that monitored synaptic outputs are produced by mixtures of inhibitory inputs and are influenced by disinhibition. For paired recordings, we stereotactically injected at P21 the mPFC of *Nrxn123* cKO/Pv-Cre and control Pv-Cre mice with AAVs expressing double-floxed eYFP and analyzed the mice at P35-P40. Since we needed to use Cre-dependent labeling of Pv^+^ interneurons in order to perform paired recordings, recordings could not be obtained from littermate mice in these and the related experiments with SST^+^ interneurons described below. Therefore, in these experiments we used control mice that were generated from *Nrxn123* cKO/Pv-Cre mice by outcrossing *Nrxn123* cKO/Pv-Cre mice with Pv-Cre mice of a similar hybrid genetic background, and analyzed control Pv-Cre mice that were genetically separated 3 generations or less from *Nrxn123* cKO/Pv-Cre mice, suggesting that they harbor a similar genetic background. Only age-matched control and test mice were used, which were manipulated in parallel at the same time.

We patched adjacent Pv^+^ interneurons and pyramidal neurons and confirmed the fast-spiking properties of the Pv^+^ interneurons by current injections (Figure 3F, G). We probed for synaptic connections by monitoring monosynaptic IPSCs (Figure 3H). Slices from *Nrxn123* cKO/Pv-Cre and control Pv-Cre mice exhibited a similar rate of success in establishing connected pairs during paired recordings (32 successes/88 pairs tested vs. 52 successes/141 pairs tested, respectively; see Table 1). However, the IPSC amplitudes in Pv^+^ interneuron pyramidal neuron synapses were decreased by ∼50% in *Nrxn123* cKO/Pv-Cre mice compared to control Pv-Cre mice (Figure 3I), similar to the decrease in the synaptic vGAT signal (Figure 3B). In neurexin-deficient Pv^+^ neurons, high-amplitude synaptic responses were preferentially lost, without a change in IPSC kinetics (Figure 3J). We detected no significant change in paired-pulse ratios, suggesting that the decrease in IPSC amplitude is likely not due to a change in release probability (Figure 3K, L). Moreover, we found no detectable change in IPSC kinetics (Table 1). Overall, these experiments indicate that the pan-neurexin deletion from Pv^+^ interneurons severely impairs the formation of synaptic connections.

**Table 1.** Properties of synapses formed by Pv^+^ or SST^+^ interneurons on pyramidal neurons in layer 5 of the mPFC as revealed by paired recordings.

| Properties | $Pv^+$ control<br>(n = 26/6) | $Pv^+$ *Nrxn123* cKO<br>(n = 46/11) | $SST^+$ control<br>(n = 10/5) | $SST^+$ *Nrxn123* KO<br>(n = 7/4) |
| --- | --- | --- | --- | --- |
| Connection probability | 36.3%<br>(32/88 tested) | 36.8%<br>(52/141 pairs) | 25.6%<br>(11/43 pairs) | 11.5%<br>(7/61 pairs) |
| Failure rate | $3.0 \pm 1.7\%$ | $3.2 \pm 1.7\%$ | $30.5 \pm 7.6\%$ | $27.1 \pm 8.0\%$ |
| Latency (ms) | $0.96 \pm 0.04$ | $1.06 \pm 0.06$ | $1.58 \pm 0.09$ | $2.23 \pm 0.11$ (***) |
| SD of latency (ms) | $0.19 \pm 0.04$ | $0.28 \pm 0.04$ | $0.51 \pm 0.05$ | $0.66 \pm 0.07$ |
| Amplitude (pA) | $269.3 \pm 50.7$ | $157.6 \pm 22.1$ (*) | $27.7 \pm 3.8$ | $17.1 \pm 0.8$ (*) |
| CV of amplitude | $0.27 \pm 0.02$ | $0.34 \pm 0.02$ | $0.33 \pm 0.02$ | $0.34 \pm 0.04$ |
| Rise time (ms) | $0.84 \pm 0.07$ | $0.85 \pm 0.06$ | $1.83 \pm 0.22$ | $2.80 \pm 0.24$(*) |
| Decay time (ms) | $18.5 \pm 0.9$ | $18.9 \pm 0.8$ | $17.0 \pm 1.7$ | $20.1 \pm 2.2$ |

Student's t-test was performed between control and cKO pairs for $PV^+$ and $SST^+$ cells, respectively (\* = $P < 0.05$ , \*\*\* = $P < 0.001$ ); number of connected pairs/mice analyzed are listed on top.

### Action potential-induced Ca^2+^ influx into the presynaptic terminals of Pv^+^ interneurons in the mPFC is not impaired by the *Nrxn123* triple deletion

To further characterize the effect of the *Nrxn123* deletion on Pv^+^ interneuron synapses, we examined their action-potential induced Ca^2+^-influx. We stereotactically injected the mPFC with AAVs expressing double-floxed mCherry at P21 and analyzed the mice at P35-P40. Pv^+^ interneurons, identified in acute slices from *Nrxn123* cKO/Pv-Cre and wild-type Pv-Cre mice by expression of mCherry, were then filled via a patch pipette with Alexa Fluor 594 (50 μM) to visualize the axonal arbor and with Fluo-5F (200 μM) to monitor presynaptic Ca^2+^-transients.

Analyses by two-photon microscopy of the density of presynaptic boutons in the axonal arbors of Pv^+^ interneurons showed that the *Nrxn123* deletion caused a modest but significant decrease in bouton density (∼15%) consistent with a decrease in synapse numbers (Figure 4A, B). We confirmed by test-current injections that the patched neurons were indeed fast-spiking neurons as expected for Pv^+^ interneurons (Figure 4C) and measured presynaptic Ca^2+^-transients as a function of presynaptic action potentials. Using this experimental configuration, we could reliably detect Ca^2+^-transients in response to single action potentials (Figure 4D). In control neurons, we observed a nearly linear relationship between the number of action potentials and Ca^2+^-transients in presynaptic nerve terminals of Pv^+^ neurons, as measured via the Ca^2+^-indicator signal (Figure 4E). Notably, the presynaptic deletion of neurexins caused no change in the amplitude or the decay kinetics of the Ca^2+^-transients, consistent with the lack of a change in paired-pulse ratio.

**Figure 4.**
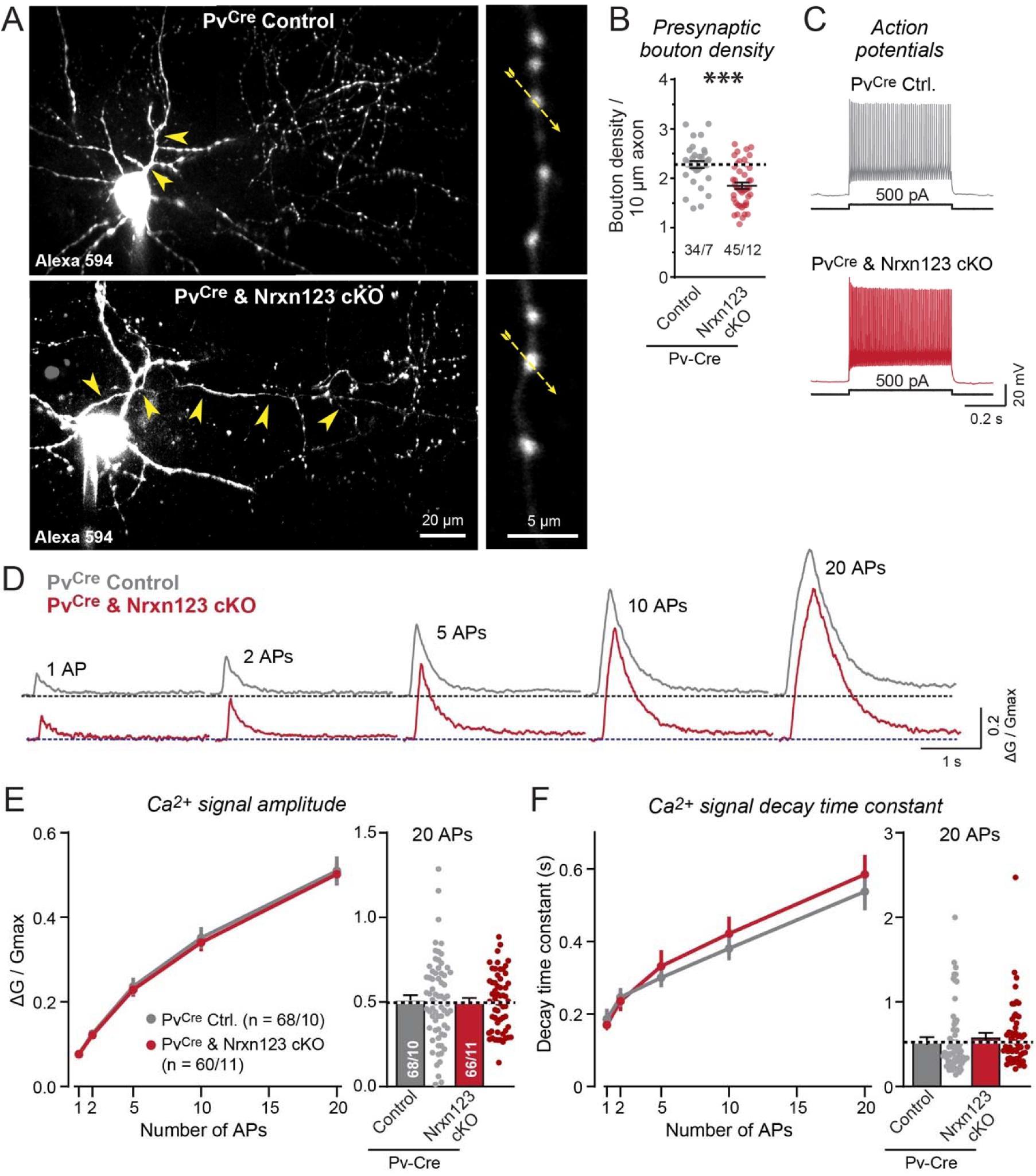
Two-photon microscopy demonstrates that pan-neurexin deletions in Pv^+^ interneurons decrease the density of presynaptic boutons in the mPFC without altering action potential-induced Ca^2+^-influx into presynaptic terminals. (A) Pv^+^ interneurons visualized by AlexaFluor 594 fluorescence in control and *Nrxn123* cKO Pv-Cre mice. Neurons identified via virally administered double-floxed mCherry were filled via the patch pipette with AlexaFluor 594 (50 μM) and Fluo-5F (200 μM; to monitor presynaptic Ca^2+^-transients) (left, representative z-stack projections of an entire neuron (arrowheads = axon); right, zoomed-in axonal boutons (dashed arrows = line scans during action potential stimulation)). (B) Summary graph of axonal bouton densities (circles, individual axons; lines, means ± SEM). (C) Representative traces of fast-spiking Pv^+^ interneuron firing. (D) Representative traces of Ca^2+^-transients monitored via Fluo-5F fluorescence in presynaptic boutons by line scans. Trains of 1, 2, 5,10, 20 APs were elicited at 50 Hz by current pulse injections (2 nA, 1 ms) through recording pipettes. (E) Summary plots of the peak amplitudes (△G/Gmax) of presynaptic Ca^2+^-transients as a function of action potential numbers (right, summary of Ca^2+^-transients peak amplitudes evoked by 20 APs). (F) Same as (**E**), but for the decay times (time of 100% to 37% amplitude decay). Data in B and E are means ± SEM (numbers indicate number of axon branches/neurons analyzed [B] or boutons/neurons analyzed [E]); statistical comparisons were performed with Student’s t-test (***p<0.001) or two-way ANOVA followed by Bonferroni *post-hoc* test; non-significant comparisons are not labeled.

Viewed together, these results suggest that deletion of presynaptic neurexins from Pv^+^ interneurons impairs the assembly of synapses formed by Pv^+^ neurons on pyramidal neurons and decreases the strength of these synapses without changing their action potential-induced Ca^2+^-influx.

### The *Nrxn123* deletion does not significantly alter inhibitory synapse numbers formed by SST^+^ interneurons in the cerebellum, hippocampus or mPFC

To test whether neurexins perform a similar function in another type of inhibitory synapse that differs from those of Pv^+^ interneurons, we analyzed synapses formed by SST^+^ interneurons. We used the same strategy as outlined above for Pv^+^ interneurons, except that we employed SST-Cre mice instead of Pv-Cre mice (Figure S1B5). Although SST is expressed in the cerebellum during early postnatal development (60), we detected no change in the density of vGluT1^+^, vGluT2^+^, or vGAT^+^ puncta in cerebellar sections, suggesting that the pan-neurexin deletion in SST^+^ cerebellar neurons does not cause a major change in excitatory or inhibitory synapse densities (Figure S6). A similar observation was made for the hippocampal CA1 region, where the pan-neurexin deletion in SST^+^ neurons again caused no detectable change in the density of vGluT1^+^ or vGAT^+^ puncta (Figure S7A-F). Moreover, when we measured the frequency and amplitude of miniature EPSCs (mEPSCs) and mIPSCs in acute slices from the CA1 region, we detected no significant change as a result of the pan-neurexin deletion in SST^+^ neurons (Figure S7G-L).

Next, we monitored synapses in the mPFC as a function of the *Nrxn123* deletion in SST^+^ neurons (Figure 5). Again, we detected no significant change in excitatory vGluT1^+^ and in inhibitory vGAT^+^ puncta densities, although there was a trend towards a decrease for the latter (Figure 5A-C). We then measured the mIPSC frequency and amplitude in layer 5 pyramidal neurons and also detected no significant mIPSC change in pyramidal neurons after the *Nrxn123* deletion in SST^+^ neurons (Figure 5D-F).

**Figure 5.**
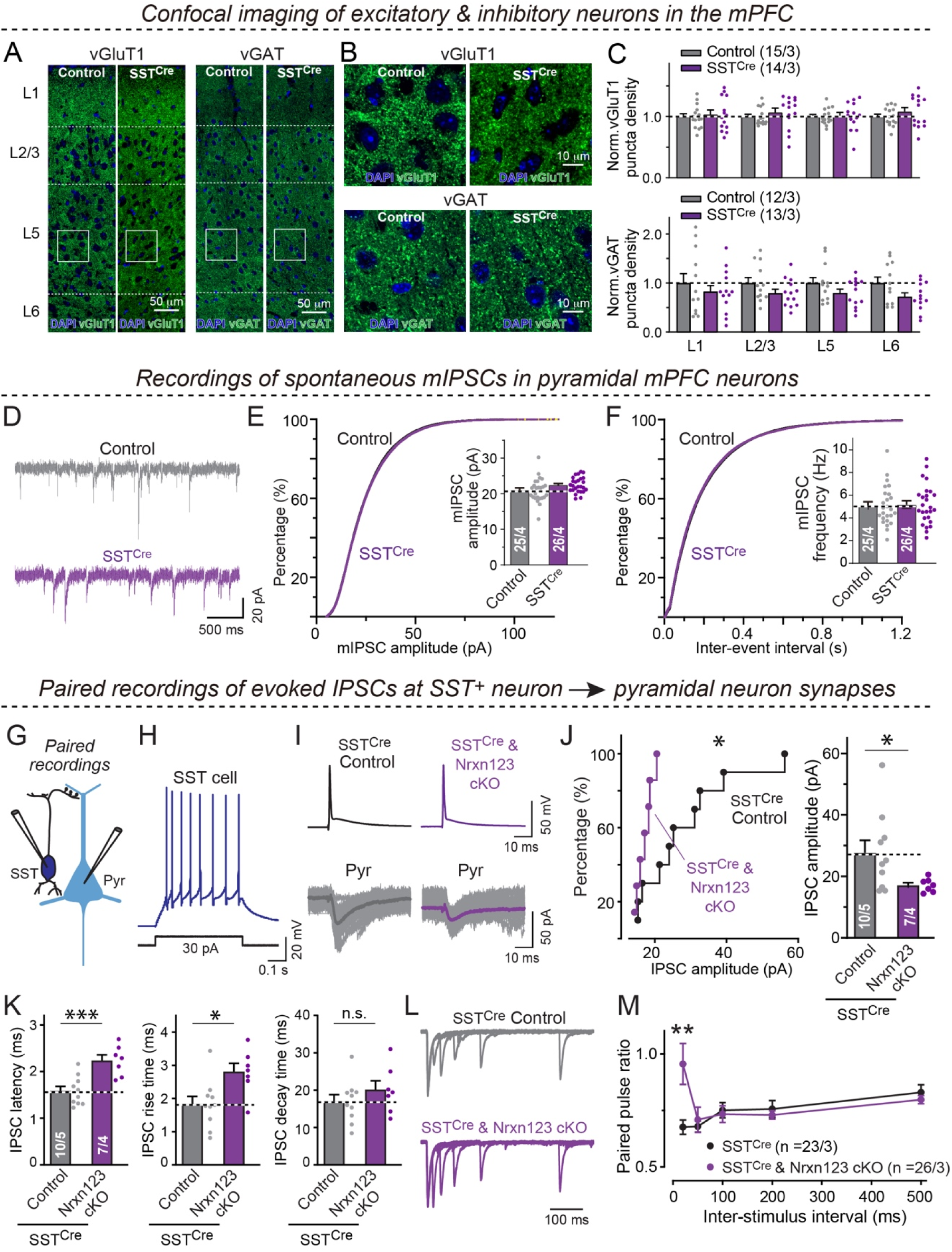
Pan-neurexin deletions in SST^+^ interneurons of the mPFC robustly impair evoked synaptic transmission as monitored by paired recordings withing significantly decreasing inhibitory synapse numbers. **(A-C)** The pan-neurexin deletion in SST⁺ interneurons does not significantly affect the density of vGluT1- and vGAT-positive synaptic puncta in the mPFC (A, representative confocal overviews (L1-L6, cortical layers); B, higher-magnification views of the boxed regions in A; C, Summary graphs of vGluT1⁺ (top) and vGAT+ puncta density (bottom) for each cortical layer). (**D-F**) The pan-neurexin deletion in SST⁺ interneurons has no effect on mIPSCs recorded from layer 5 pyramidal neurons in mPFC (D, Representative mIPSC traces; E & F, Cumulative distributions of mIPSC amplitudes and inter-event intervals, respectively, with summary graphs of the mean mIPSC amplitude and frequency). (G) Paired recording configuration to monitor inhibitory synapses formed by SST^+^ interneurons on distal dendrites of pyramidal neurons (Pyr). (H) Representative traces of a low-threshold firing pattern of SST^+^ interneurons. (I) Representative paired-recording traces showing unitary synaptic connections between SST^+^ interneurons and pyramidal neurons. IPSCs were evoked by current injections into patched presynaptic SST^+^ interneurons (top), and recorded from patched postsynaptic pyramidal neurons (bottom grey traces, individual responses; black/purple traces, averaged responses). (J) Cumulative probability plot of unitary mean IPSC amplitudes (inset = summary graph of IPSC amplitudes). (K) Summary plots of the latency (left), rise time (center), and decay time (right) of unitary IPSCs in SST-interneuron pyramidal neuron synapses. **(L** & **M)** Representative traces showing PPRs of unitary IPSCs in SST-interneuron pyramidal neuron synapses at increasing inter-stimulus intervals (L), and summary graph of the mean PPRs as a function of the inter-stimulus interval (M). Unitary IPSCs were evoked by optogenetic stimulation of SST^+^ cells after stereotactic injection of the mice with AAVs encoding CAG-DIO-ChR2-tdTomato. Data are means ± SEM; statistical comparisons were performed with Student’s t-test (*p<0.05; **p<0.01; ***p<0.001). Numbers indicate the number of cells and mice examined.

### The *Nrxn123* deletion significantly decreases the presynaptic release probability in SST^+^ interneurons

Although the experiments described above detected no overall phenotype in inhibitory synapses that was induced by the pan-neurexin deletion in SST^+^ neurons, a phenotype could have been missed since only a subset of inhibitory neurons express SST and this subset might also be contributing to disinhibitory circuits (61). Thus, we performed paired recordings, comparing *Nrxn123* cKO/SST-Cre with wild-type SST-Cre mice (Figure 5G-I). Similar to the paired recordings of Pv^+^ wild-type and *Nrxn123* cKO neurons, we detected no significant difference between the connection frequency of SST^+^ wild-type and *Nrxn123* cKO neurons, although there was a trend towards fewer pairs in the *Nrxn123* cKO neurons (11 successes/43 pairs vs. 7 successes/61 pairs, respectively; Table 1). Strikingly, we found that the presynaptic deletion of neurexins in SST^+^ neurons caused a large decrease in the IPSC amplitude (∼40%; Figure 5J). No SST^+^ interneuron pyramidal neuron synaptic connection exhibited an IPSC amplitude of >20 pA in *Nrxn123* cKO/SST-Cre mice, whereas >50% of these synaptic connections exhibited IPSCs with amplitudes of >20 pA in wild-type mice (Figure 5J). In addition, IPSCs elicited by stimulation of neurexin-deficient SST^+^ interneurons were slower, as indicated by an increased synaptic latency (∼30%) and increased rise time (∼30%) (Figure 5K; Table 1).

To test whether the decrease in synaptic strength in SST^+^ interneuron synapses is due to a decrease in release probability, we measured the PPR at SST^+^ interneuron pyramidal neuron synapses. For this purpose, the low synaptic strength of individual connections made it necessary to simultaneously stimulate multiple synaptic inputs from SST^+^ interneurons onto the same postsynaptic neuron. We achieved this by stereotactically infecting the mPFC of Nrxn123 cKO/SST-Cre and wild-type SST-Cre mice at P21 with AAVs that encode a double-floxed expression cassette for a fusion protein composed of channelrhodopsin-2 and tdTomato (62). We then recorded IPSCs evoked by light stimulation from pyramidal neurons in layer 5 of the mPFC at P35-P40. The stimulation intensity was adjusted until a typical monosynaptic response with linear rise and exponential decay times was obtained and an initial IPSC amplitude of 0.1-0.2 nA. Although this experimental setup does not allow measurements of synaptic strength, it enables the administration of two closely spaced stimuli at defined intervals to monitor relative IPSC amplitudes during paired-pulse ratio measurements (Figure 5L). Using this approach, we observed a large increase in paired-pulse ratios selective for short interstimulus intervals (Figure 5M). Together with the slower IPSC kinetics, these findings indicate that pan-neurexin deletion in SST^+^ interneurons reduces the presynaptic release probability.

### The *Nrxn123* deletion in SST^+^ interneurons significantly impairs action potential-induced Ca^2+^ influx

The decrease in release probability that is induced by the neurexin deletion in synapses formed by SST^+^ interneurons in the cortex resembles the phenotype of the pan-neurexin deletion in the calyx of Held synapse that is caused by a decrease in action potential-triggered Ca^2+^-influx (49). Thus, we tested whether action potential-induced Ca^2+^-influx is decreased by the *Nrxn123* deletion in SST^+^ interneurons (Figure 6), using the same approach as described above for Pv^+^ interneurons (Figure 4). Consistent with the lack of a change in synapse density, we detected no change in the density of presynaptic boutons as a result of the neurexin deletion (Figure 6A, B). SST^+^ interneurons from *Nrxn123* cKO/SST-Cre and SST-Cre control mice exhibited a typical low-threshold spike firing pattern (Figure 6C). Ca^2+^-transients induced by single action potentials could be reliably identified in individual presynaptic boutons (Figure 6D).

**Figure 6.**
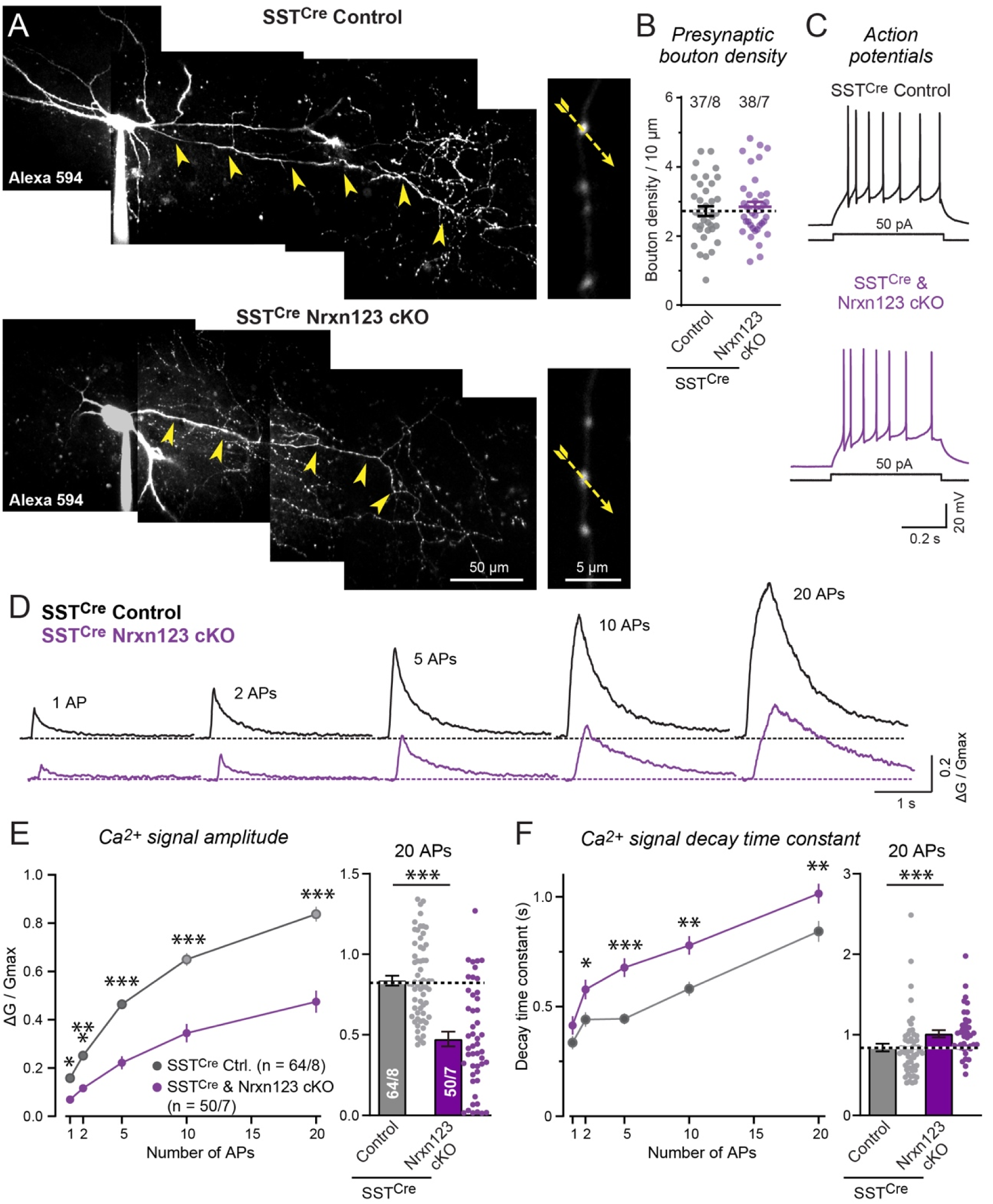
Two-photon microscopy reveals that the pan-neurexin deletion in SST^+^ interneurons in the mPFC suppresses action potential-evoked presynaptic Ca^2+^-transients. (A) Pv^+^ interneurons visualized by AlexaFluor 594 fluorescence in control and *Nrxn123* cKO SST-Cre mice. Neurons identified via virally administered double-floxed mCherry were filled via the patch pipette with AlexaFluor 594 (50 μM) and Fluo-5F (200 μM; to monitor presynaptic Ca^2+^-transients) (left, representative z-stack projections of an entire neuron (arrowheads = axon); right, zoomed-in axonal boutons (dashed arrows = line scans during action potential stimulation)). (B) Summary graph of axonal bouton densities (circles, individual axons; lines, means ± SEM). (C) Representative traces of low-threshold firing in SST^+^ interneurons. (D) Representative traces of Ca^2+^-transients monitored via Fluo-5F fluorescence in presynaptic boutons by line scans. Trains of 1, 2, 5,10, 20 APs were elicited at 50 Hz by current pulse injections (2 nA, 1 ms) through recording pipettes. (E) Summary plots of the peak amplitudes (△G/Gmax) of presynaptic Ca^2+^-transients as a function of action potential numbers (right, summary of Ca^2+^-transients peak amplitudes evoked by 20 Aps). (F) Same as (**E**), but for the decay times (time of 100% to 37% amplitude decay). Data are means ± SEMs (numbers indicate number of axon branches/neurons analyzed [B] or boutons/neurons analyzed [E]); statistical evaluations were performed by Student’s t-test or two-way ANOVA with Bonferroni *post-hoc* tests (* = p<0.05; *** = p<0.001).

Importantly, we observed that the neurexin deletion produced a large decrease in the amplitude of action-potential-induced Ca^2+^-transients (Figure 6E). In control SST^+^ neurons, the Ca^2+^-transient increased almost linearly as a function of action potential numbers. In neurexin-deficient SST^+^ interneurons, the Ca^2+^-transient amplitude was uniformly decreased by ∼50%, and the Ca^2+^-transient decay time was significantly increased (Figure 6D-F). These changes explain the decrease in release probability, suggesting that the neurexin deletion suppresses synaptic strength in SST^+^ interneuron synapses at least in part by lowering Ca^2+^-influx.

### Pv^+^ and SST^+^ interneurons do not exhibit major differences in the expression of neurexins and other synaptic cell-adhesion molecules as analyzed by single-cell quantitative RT-PCR

A possible explanation for the dramatically different phenotypes of the pan-neurexin deletions in Pv^+^ vs. SST^+^ interneurons of the mPFC is that these neurons might selectively express different neurexins. To test this hypothesis, we compared the relative expression profiles of neurexins in inhibitory Pv^+^ and SST^+^ interneurons of the mPFC using Fluidigm-based quantitative single-cell RT-PCR of cytoplasm aspirated from the neuronal cell bodies via a patch pipette. Specifically, we identified Pv^+^ and SST^+^ interneurons in acute slices from wild-type Pv-Cre or SST-Cre mice injected with AAVs expressing double-floxed eYFP at P21 and analyzed at P35-P40. We patched the labeled neurons, aspirated cytosol, and performed mRNA measurements on the cytosol (Figure 7). We found that apart from typical markers for Pv^+^ and SST^+^ interneurons (such as parvalbumin, somatostatin, and synaptotagmin-2 (63)), both types of interneurons expressed similar overall levels of neurexins and of other synaptic cell-adhesion molecules (Figure 7). Lphn1 and Slitrk3 exhibited modest differences between the two types of interneurons, but all three neurexin genes were expressed similarly. Given the small number of synaptic adhesion molecules surveyed here, our results do not exclude the likelihood that the distinct functions of neurexins in Pv^+^ and SST^+^ neurons are conditioned by the differential expression of synaptic adhesion molecules that remain to be identified.

**Figure 7.**
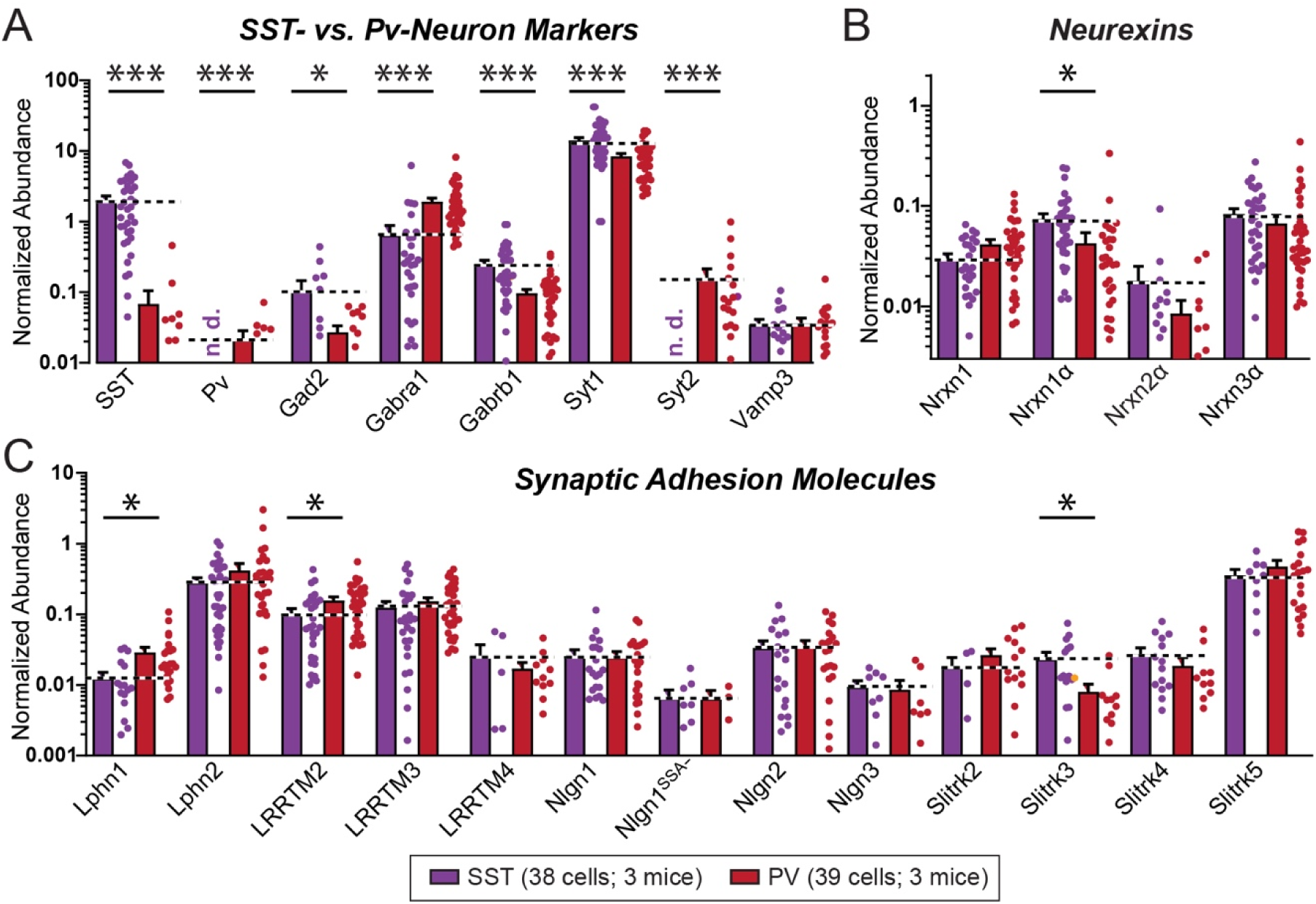
Quantification of mRNA expression levels in patched Pv^+^ or SST^+^ interneurons from the mPFC fails to identify dramatic differences in the expression of neurexins or other synaptic adhesion molecules. **(A)** Average single-cell expression levels of the indicated markers in Pv^+^- and SST^+^- interneurons, measured by qRT-PCR in cytosol from patched neurons as described (45). **(B** & **C**) Same as A, but for mRNAs encoding the three principal α-neurexins (B) or synaptic adhesion molecules other than neurexins (C). Data are means ± SEM (n = 38/3 and 39/3 cells/mice for SST^+^ and Pv^+^ neurons, respectively). Statistical evaluations were performed by Student’s t-test with * = p<0.05; *** = p<0.001 since each qRT-PCR assay is a separate experiment.

## Discussion

A large number of studies attest to the physiological importance of neurexins, showing that mutations or deletions of neurexins in organisms ranging from worms to humans cause synaptic dysfunction (37–41). Moreover, alterations in neurexin genes, in particular *NRXN1*, are associated with diverse neuropsychiatric disorders, including autism, schizophrenia, and Tourette syndrome, and anti-neurexin antibodies are prominent agents in autoimmune encephalitis (64–69). Thus, with thousands of papers listed in PubMed, neurexins are deeply researched. Nevertheless, considerable uncertainty exists about the overall functions of neurexins. The physiological consequences of neurexin mutations differ dramatically between experimental preparations and organisms, including the original observation that deletion of all α-neurexins severely impairs mouse survival and synaptic transmission (11) and more recent findings that *Nrxn3* deletions decrease the presynaptic release probability at inhibitory synapses in olfactory bulb neurons (20, 27) and that alternative splicing of *Nrxn1* and *Nrxn3* at splice-site 4 controls postsynaptic NMDARs and AMPARs, respectively, at excitatory synapses (19, 23–25).

A decade ago, we published a study aiming to compare fundamental neurexin functions between diverse synapses by conditionally deleting all neurexins except for Nrxn1γ in different types of neurons (47). This pivotal study established that pan-neurexin deletions produced distinct phenotypes in different synapses, a result that was amplified in subsequent papers by examining other types of neurons with similar approaches (49, 50, 52–55). However, we recently retracted the foundational Chen et al. (2017) paper (47, 48) because AI-based image analyses discovered microduplications of image areas in three panels of the paper, even though these image abnormalities had no significant impact on the paper’s conclusions. As a result, the finding of distinct neurexin functions in diverse synapses was removed from the scientific community, motivating the present study. Here, using a complete reanalysis of existing raw data from the original study and newly performed complementary experiments, we affirm and expand the original paper’s results. Moreover, to ensure that future AI-based analyses do not detect any anomalies in published images, we are making all raw data publicly available (SDR: https://purl.stanford.edu/xp149zq3345). Our present findings confirm, with additions and alterations, the original paper’s findings and extend them to additional synapses, validating the overall conclusion that neurexins are universal regulators of synapse assembly that differentially impact diverse synapses.

Our results establish two major specific observations that motivate the overall conclusion of diverse, neuron type-specific neurexin functions. First, in excitatory climbing-fiber synapses and inhibitory hippocampal cortical Pv^+^ neuron pyramidal neuron synapses, the pan-neurexin deletion produces a robust impairment in synapse assembly, leading to synapse loss (Figure 1-4; S2-5). Most illuminating here are our paired recordings in the mPFC that revealed that the pan-neurexin deletion reduced the IPSC amplitudes of Pv^+^ interneuron pyramidal neuron synapses by ∼50% (Figure 3I) and decreased the density of synaptic boutons measured in the two photon imaging experiments by ∼20% (Figure 4B) without affecting the Ca^2+^-dynamics of presynaptic Pv^+^ nerve terminals (Figure 4D-F). Thus, the pan-neurexin deletion likely primarily impairs synapse assembly by rendering synapses smaller and weaker, with a secondary loss of synapses. We observed a similar loss of synapses in the hippocampus but could not detect a change in synaptic transmission here, probably because we measured all synaptic responses in slices using extracellular stimulation instead of paired recordings (Figure S4).

Second, the pan-neurexin deletion severely impaired the function of synapses formed by SST^+^ interneurons without significantly changing synapse numbers (Figures 5, 6). The functional impairment of SST^+^ interneuron synapses supports an essential role for neurexins in enabling the localization of Ca^2+^-channels at synaptic release sites, as described originally for the triple α-neurexin KO mice (11) and previously also confirmed for pan-neurexin KO mice in brainstem synapses (49, 50). This phenotype does not necessarily imply a direct role of neurexins in Ca^2+^-channel recruitment, a role performed by RIMs and RIM-BPs (70, 71), but could reflect an indirect effect. It is striking that none of the pan-neurexin KO phenotypes described here seems similar to the neurexin-3 cKO phenotypes we described in other inhibitory synapses (20, 27), suggesting that we might discover an even larger range of phenotypes in other types of synapses.

Multiple questions arise at this point. 1) How can we explain the differences in phenotypes observed in different types of synapses? A plausible hypothesis is that the relative abundance of different neurexins, their various splice variants, and their multifarious pre- and postsynaptic ligands, and not the absolute amounts of these proteins, dictate synaptic functions (45). Although we detected no major differences in the expression of neurexins between Pv^+^ and SST^+^ neurons (Figure 7), differences in alternative splicing of neurexins have large effects on at least some synapse properties (19, 23–25, 27). As a result, presynaptic neurexins might mediate distinct functions even in synapses formed by the same presynaptic neurons onto different postsynaptic cells, depending on the ligands that are expressed of these cells. This may be a general principle of synaptic adhesion molecules. Alternatively, it is conceivable that canonical neurexin functions are differentially compensated in various types of synapses by neurexin-independent pathways. The chronic nature of our cortical manipulations could give time for distinct compensatory processes in different types of neurons. As another similarly unlikely possibility given the known properties and localization of neurexins, it is conceivable that the pan-neurexin deletions cause distinct phenotypes in different types of neurons because they impair a common upstream process that is transformed into distinct downstream consequences in various types of synapses. Future experiments studying specific neurexin functions in depth will have to address these questions.

1. Why isn’t the phenotype more severe? In some analyses of single neurexins or neurexin splice variants, more extensive phenotypes were found than what we describe here. For example, constitutive expression of *Nrxn3*^SS4+^ in CA1 region neurons causes a large (>50%) decrease in postsynaptic AMPARs at CA1 subiculum synapses (19), whereas deletion of *Nrxn3* from inhibitory granule cells in the olfactory bulb greatly decreases (>50%) the presynaptic release probability (27). A plausible reason for this difference in phenotype severity is that Cre-mediated recombination of multiple neurexin genes may not be 100%, leading to an underestimate of the phenotype. Moreover, the phenotypes we detect clearly depend on the synapses analyzed since we did not study CA1 region output synapses or olfactory granule cell output synapses in the present study. Although the phenotypes observed in the paired recordings in the cortex are very robust (Figures 3-6), we may have, by chance, examined synapses with weaker phenotypes. Finally, it may be the ratio of different neurexins more than their absolute levels that determine phenotypes, resulting in an occlusion of phenotypes when multiple neurexins are deleted.
2. Why do morphological and electrophysiological phenotypes sometimes differ in severity? This is observed, for example, for Pv^+^ synapses in the hippocampus (Figure S4). One explanation may be that some synapses could undergo a homeostatic adjustment that renders them stronger, compensating for the loss of synapses. Another explanation could be that the functional analysis in this example is less specific than the paired recordings performed for Pv^+^ and SST^+^ synapses in the cortex, where the resulting functional and imaging phenotypes are similarly severe (Figures 3-6). A technical limitation of all synapse density measurements by immunolabeling is that the apparent magnitude of a phenotype is inherently sensitive to the post-acquisition image analyses (Figure S9). As a consequence, the results obtained by morphological analyses, such as synapse density quantifications, depend critically on the intensity and object-sizing thresholds chosen to differentiate signal from background noise

Finally, our study includes inherent and important limitations. We do not know whether the Cre-driver mice and the virally delivered Cre mediated full recombination of the *Nrxn1*, *Nrxn2*, and *Nrxn3* genes in our experiments. This question is difficult to address experimentally because the targeted cells in our experiments mostly involve a small subset of neurons and sensitive specific antibodies for neurexins are not readily available. We measured the levels of neurexin mRNAs in the inferior olive after AAV-mediated neurexin deletions and did observe an incomplete deletion of neurexins (Figure 1B), but these measurements were not definitive because the deletion targeted neurons whereas glia also express neurexins. In addition, incomplete nonsense-mediated mRNA decay may have retained non-functional mRNAs. The fact that in a previous study (72) the sextuple KO of PTPRs and neurexins causes a massive phenotype indicates that Pv-Cre-mediated recombination is effective in at least some neuron types but proving the extent of this recombination is technically challenging.

Moreover, the *Nrxn1* deletion we analyzed does not abolish expression of *Nrxn1γ*, which does not bind to any of the known neurexin ligands except for FAM19A’s and CA10/11 (6, 73). In C. elegans, which expresses a single α- and γ-neurexin but lacks β-neurexin, the γ-neurexin is functionally important (7). A recent study also indicated a major function for *Nrxn1γ* in mouse neurons that was ascribed to its heparan sulfate modification (74). Thus, it is possible that some of the functions of neurexins were not detected in our analysis because the expression of *Nrxn1*γ persists.

Another limitation derives from the fact that paralogs of genes do not necessarily perform comparable functions. The three α- and β-neurexin genes are highly homologous and bind to the same ligands but, when analyzed in the same synapses, appear to have distinct functions (23, 24). The key here is the fact that nearly all neurons co-express all seven principal neurexins (the α-, β-, and γ-neurexins) but in different ratios and splice variants. Deleting individual neurexins causes distinct phenotypes whenever studied in detail. As a result, it is possible that the double and triple neurexin deletions produce not simply the additive or even synergistic effects of single deletions, but may also cause unpredictable functional interactions that might compensate for phenotypes.

### Summary

Consistent with earlier results, our data indicate that neurexins perform a limited essential role in establishing synapse formation but are essential for shaping synapses in a manner that depends on the identity of the pre- and postsynaptic neuron. In SST^+^ interneuron synapses, neurexins couple Ca^2+^-channels to presynaptic release sites as shown earlier for other synapses (11, 49). In cortical Pv^+^ interneuron synapses, neurexins are required for assembly of functional synapses but not for coupling Ca^2+^- channels to presynaptic release sites. The phenotypes are severe in each type of synapse, suggesting that neurexins are organizers and not modulators of synapses. Moreover, the phenotypes are too different between different types of synapses to be attributed to a similar basic mechanism. A molecular dissection of the function of each neurexin in different types of synapses will likely be required to understand why their functions are different, which will then allow relating the particular molecular mechanism involved to the overall design of synapses. In general, however, it is clear that neurexins do not perform just one unitary function, with the lack of apparent roles for neurexins overall in some synapses possibly due to a mixture of functions that balance each other. Thus, the key conclusion of our study comparing defined synapses in three brain regions is that neurexins as a whole perform distinct functions in different types of synapses and brain regions.

## Materials and Methods

Detailed information on all protocols, reagents, antibodies, software, and other resources used in this study is provided in the SI Appendix.

### Experimental model: *Nrxn123* triple cKO mice

The use and care of animals complied with the guidelines of the Administrative Panel on Laboratory Animal Care at Stanford University and the Animal Care and Use Committee of Huazhong University of Science and Technology, China. All mouse strains, genetic crosses, and primers used for genotyping are described in the SI Appendix, Extended Methods.

### Virus productions and stereotactic *in vivo* injections

Lentiviruses and AAVs were produced as described (56, 75, 76). For inferior olive manipulations, lentiviruses expressing EF1a-NLS-HA-ΔCre-P2A-tdTomato-T2A-EGFP::Syb2 or EF1a-NLS-HA-Cre-P2A-tdTomato-T2A-EGFP::Syb2, or AAVs expressing hSyn-GFP-Cre or hSyn-GFP-ΔCre were stereotactically injected at P1 as described (77). For mPFC manipulations, AAVs expressing Ef1α-DIO-eYFP (AAV-DJ), Ef1α-DIO-mCherry (AAV5) or CAG-DIO-ChR2-tdTomato (AAV-DJ) were stereotactically injected at P21 as described (78). See detailed Extended Methods in the SI Appendix.

### Electrophysiology

Slices (300 μm thick) were prepared using a vibratome (LEICA VT1200S) as described previously (79). For climbing-fiber (CF)-EPSC recordings (Figure 2G-M and Figure S3), sagittal slices (300 μm thick) of the cerebellum were prepared from mice at postnatal day 35 (P35). Picrotoxin (50 μM) and CNQX (2 μM) were added to the extracellular solution during whole-cell recordings. For the recordings of inhibitory synaptic transmission in the CA1 (Figure S4C-N), horizontal slices (300 μm thick) of the hippocampus were prepared from mice at approximately P45-55. D-APV (50 μM) and CNQX (20 μM) were added to the extracellular solution during whole-cell recordings. Coronal mPFC slices were cut at P35-40. Paired recordings (Figure 3F-J and 5G-K) were recorded in Pv^+^ or SST^+^ interneurons (in current-clamp mode) and nearby connected pyramidal (Pyr) cells. Miniature IPSCs (mIPSCs) were recorded in pyramidal cells in the presence of 1 μM TTX, 20 μM CNQX, and 50 μM D-APV. See detailed methods in SI Appendix, Extended Methods.

### Two-photon Ca^2+^ imaging

For Figures 4 and 6, we utilized Fluo-5F (200 μM) as a Ca^2+^ indicator to monitor Ca^2+^ changes. Alexa Fluor 594 (50 μM) was added into the internal solution for visualization of the cell morphology. For details, see Extended Methods in the SI Appendix.

### Immunohistochemistry

Immunohistochemical experiments were performed as previously described (72). Primary antibodies (guinea pig anti-vGluT1, 1:1000; guinea pig anti-vGluT2, 1:500; mouse anti-Calbindin, 1:500; rabbit anti-Calbindin, 1:500; rabbit anti-GFP, 1:1000; rabbit anti-vGAT, 1:500; rabbit anti-Pv, 1:2000) and secondary antibodies (1:1000, Invitrogen, Alexa 488, Alexa 546, or Alexa 633) were used for immunolabeling. See detailed methods in SI Appendix, Extended Methods.

### Confocal image acquisition and analysis

Images (at 1024 × 1024 resolution) were acquired using a Nikon confocal microscope (A1Rsi) with a 60X oil objective (PlanApo, NA1.4). The general analysis module in NIS-Elements Advanced Research software (Nikon) or ImageJ software was used. See detailed methods in SI Appendix, Extended Methods.

### STED Microscopy

STED imaging was performed as previously described (27). Immunolabeling was performed using guinea pig anti-vGluT2 (1:500) as the primary antibody, followed by incubation with STAR ORANGE-conjugated secondary antibody (1:500). See detailed methods in SI Appendix, Extended Methods.

### qRT-PCR and single-cell transcriptional profiling

For Figure 1B, qRT-PCR was performed as previously described (27). The following predesigned assays were used (gene, assay ID): Actb (Mm.PT.39a.22214843.g), Nrxn1 (Mm.PT.58.33547773), Nrxn3 (Mm.PT.58.30432445). Nrxn2 mRNAs were probed using the following assays (gene, primer 1, primer 2, probe): Nrxn2 (5’-CTATACATGGCCTCCCAATGAC-3’, 5’-CGCTGGTGTGTGCTGAA-3’, 5’-FAM/AGTACACGG/ZEN/ATGGACCGC-3’).

For Figure 7, Single-Cell Transcriptional Profiling was performed as previously described (45). Primer and probe sequences used for the single-cell analyses are listed in Table S2. See detailed methods in SI Appendix, Extended Methods.

### Quantifications and statistical analyses

All data are means ± SEM; numbers of neurons, sections, or ROIs /mice analyzed are shown in the bars or plots. Inter-group comparisons were performed using two-tailed Student’s *t*-test for comparisons of two conditions (bar diagrams) or Kolmogorov-Smirnov test for cumulative distributions, multiple comparisons were analyzed with two-way ANOVA with Bonferroni’s post-test (*p<0.05, **p<0.01, and ***p<0.001). Image backgrounds were normalized using the same settings for all conditions in an experiment, and immunoreactive elements were analyzed with the Nikon analysis software. Test and controls were analyzed on anonymized samples to prevent observer bias. All experiments were performed with at least three independent biological replicates.

## Data availability

All raw data are publicly available at (SDR: https://purl.stanford.edu/xp149zq3345)

## Acknowledgments

We thank Dr. L.Y. Chen for initiating this project 15 years ago and Drs. L. Chen, R. L. Zhong, and Y. Wu for invaluable help with Ca^2+^-imaging experiments. Early stages of this study were supported by grants from the NIMH (MH052804 to T.C.S.).

## Contact for Reagent and Resource Sharing

Requests for resources and reagents should be directed to the Lead Contact, Thomas C. Südhof

## Author Contributions

M.J., Y.L. and M.A., mouse genetics; M.J., Y.L., S. L., W.Z. and B.Z., electrophysiology; Y.L., M.A., and L.Z. morphology and immunohistochemistry; M.J., two-photon imaging; O.G. and M.J., single-cell qRT-PCR analyses; M.J., Y.L., M.A., B.Z., and T.C.S. planned the experiments, analyzed the data, and wrote the paper.

