## Supplementary material for "Conditional Deletion of All Neurexins Defines Diversity of Essential Synaptic Organizer Functions for Neurexins": SI Appendix

**This PDF file includes:**

Extended Materials and Methods

Figures S1 to S9

Tables S1 and S2

SI References

**Extended material and methods**

**Experimental model: *Nrxn123* triple cKO mice**

All use and care of animals complied with the guidelines of the Administrative Panel on Laboratory Animal Care at Stanford University and the Animal Care and Use Committee of Huazhong University of Science and Technology, China. Triple floxed *Nrxn1*, *Nrxn2*, and *Nrxn3* mice (*Nrxn123*^f/f^) were generated by flanking exon 18 according to the exon counts in ref. *5* with FloxP sites. Exon 18 is an out-of-frame exon that encodes the N-terminal part of LNS6 which is shared by all α- and β-neurexins but not by *Nrxn1γ* (Figure S1A). Pv-IRES-Cre, SST-IRES-Cre, Crh-IRES-Cre driver lines were crossed with *Nrxn123*^f/f^ mice to generate *Nrxn123* KO mice with pan-neurexin deletions in the corresponding target neurons. Mice were maintained on a mixed BL6/C57; CD1 background; mice of both sexes were used. Breeding cages were maintained by crossing Pv-Cre/+; *Nrxn123*^f/f^, Crh-Cre/+; *Nrxn123*^f/f^ or SST-Cre/+; *Nrxn123*^f/f^ with *Nrxn123*^f/f^ mice. Because it is difficult to precisely identify Pv^+^ and SST^+^ cells from *Nrxn123*^f/f^ mice, we used Pv-Cre and SST-Cre mouse lines without *Nrxn123*^f/f^ as controls in paired recording and Ca^2+^ imaging experiments. These mice were generated from Pv-Cre/+; *Nrxn123*^f/f^ or SST-Cre/+; *Nrxn123*^f/f^ mice by outcrossing them with Pv-Cre or SST-Cre mice on a similar genetic background. Subsequently, paired recording and Ca^2+^-imaging experiments analyzed age-matched test and control mice that were separated by 3 generations or less. All other experiments used littermate *Nrxn123*^f/f^ mice without Cre expression as controls. Note that *Nrxn123* cKO mice were originally reported by Chen et al. (2017) (1), deposited in Jackson Labs, and since then extensively used by our and by other laboratories for other experiments. The mouse models used in this paper are schematically described in Figure S1B.

The following primer sequences were used for genotyping:

*Nrxn1* flox:	5’ GTAGCCTGTTTACTGCAGTTCATTCC 3’

5’ CAAGCACAGGATGTAATGGCCTTTC 3’

*Nrxn2* flox:	5’ CAGGGTAGGGTGTGGAATGAGGTC 3’

5’ GTTGAGCCTCACATCCCATTTGTCT 3’

*Nrxn3* flox:	5’ AATAGCAGAGGGGTGTGACAC 3’

5’ CGTGGGGTATTTACGGATGAG 3’

Cre:	5’ GAACCTGATGGACATGTTCAGG 3’

5’ AGTGCGTTCGAACGCTAGAGCCTGT 3’

Crh-Cre:	5’ CTTACACATTTCGTCCTAGCC 3’

5’ CACGACCAGGCTGCGGCTAAC 3’

5’ CAATGTATCTTATCATGTCTGGATCC 3’

**Virus productions and stereotactic *in vivo* injections**

Lentiviruses and AAVs were produced as described (2-4). For lentiviral production, transfections of HEK 293T cells were performed using the calcium-phosphate method with Lenti-EF1a-NLS-HA-ΔCre-P2A-tdTomato-T2A-EGFP::Syb2 or Lenti-EF1a-NLS-HA-Cre-P2A-tdTomato-T2A-EGFP::Syb2 and three helper plasmids (pRSV-REV, pMDLg/ pRRE and pVSVG). 8–12 h after transfection, the medium was replaced with DMEM with 10% FBS. Then, 48 h after transfection, the cell medium was cleared by centrifuging in table-top centrifuge at 2,000g for 3 min, and filtered through a 0.45 μm PES membrane. The viral supernatant was loaded onto a 2 ml 30% sucrose cushion in PBS and centrifuged in the Thermo Fisher Scientific SureSpin 630 rotor at 19,000 rpm for 2 h. The viral pellet was resuspended in 50 μl MEM, aliquoted, and stored at −80 °C, AAV-hSyn-GFP-Cre and AAV-hSyn-GFP-ΔCre were prepared at Janelia Farm Viral core facility.

1. Inferior olive: P1 pups were anesthetized on ice for 5 min and immobilized on an ice bag with ear bars. Lentiviruses expressing EF1a-NLS-HA-ΔCre-P2A-tdTomato-T2A-EGFP::Syb2 or EF1a-NLS-HA-Cre-P2A-tdTomato-T2A-EGFP::Syb2, or AAVs expressing hSyn-GFP-Cre or AAV-hSyn-GFP-ΔCre were stereotactically injected into the inferior olive at P1 as described (5), then used for immunohistochemistry experiments at P28. Injection sites were confirmed after the experiments by sectioning the inferior olive.

2. Prefrontal cortex: AAVs expressing Ef1**α**-DIO-eYFP (AAV-DJ), Ef1**α**-DIO-mCherry (AAV5) and CAG-DIO-ChR2-tdTomato (AAV-DJ) were stereotactically injected into the mPFC at P21 (coordinates: 2.0 mm anterior to bregma; 0.4 mm lateral to midline; depth of ~2.1 mm from the dura, with 0.5 µl AAV injected at 0.2 µl/min with a microinjection pump) as described (6).

**Electrophysiology**

1. Cerebellum: Sagittal slices (300 μm thick) of the cerebellum from young mice (~P35) were sectioned by standard procedures with a vibratome (LEICA VT1200S) (7). Cutting solutions contained (in mM): 85 NaCl, 75 sucrose, 24 NaHCO_3_, 25 glucose, 2.5 KCl, 1.3 NaH_2_PO_4_, 4 MgCl_2_ and 0.5 CaCl_2_. Slices were recovered in artificial cerebrospinal fluid (ACSF) containing (in mM): 125 NaCl, 26 NaHCO_3_, 25 glucose, 2.5 KCl, 1.25 NaH_2_PO_4_, 1 MgCl_2_ and 2 CaCl_2_. (pH 7.4, aerated with 95% O_2_ / 5% CO_2_). For recordings of climbing-fiber (CF)-EPSCs (Figure 2G-M and Figure S3), picrotoxin (50 μM) and CNQX (2 μM) were added to the extracellular solution. Internal pipette solutions contained (in mM): 140 Cs-gluconate, 2 TEA, 10 HEPES, 10 Na_2_-phospho-creatine, 2 MgATP, 0.3 Na_2_GTP, 0.5 EGTA, 2 QX-314, pH 7.2. Whole-cell recordings in the voltage-clamp mode at -70 mV were made at room temperature with an Axon amplifier under visualization of neurons with an upright microscope (BX51WI, Olympus) equipped with a 60X water immersion objective (Olympus). Patch pipette resistances of 2-3 MΩ, and series resistances of <10 MΩ were comparable between genotypes and compensated by 80%-90%. CF-EPSCs were evoked using a matrix stimulating electrode placed in the granule cell layer adjacent to the recorded Purkinje cells to evoke all-or-none responses. A 20 Hz train of 30 pulses was applied to evoke a train of EPSCs, RRP and Pr were calculated as described (8, 9).

2. Hippocampus: Horizontal hippocampal slices (300 μm thick) were prepared from mice (age ~P45-60) and recovered as described above for cerebellar slices. Recordings of IPSCs (Figure S4C-N), were performed at room temperature as described above except that the ACSF bath solution contained D-APV (50 μM) and CNQX (20 μM) and the internal pipette solution was composed of (in mM): 140 CsCl, 10 HEPES, 5 Na_2_-phosphocreatine, 4 MgATP, 0.3 Na_2_GTP, 0.5 EGTA, 2 QX-314, pH 7.2. IPSCs were recorded from CA1 pyramidal neurons and evoked using a matrix stimulating electrode placed on the pyramidal cell layer. Paired-pulse ratios (PPRs) of evoked IPSCs were determined as the ratios of the second to the first EPSC amplitudes.

3. Prefrontal cortex (PFC): All experiments were performed at P35-40. Coronal mPFC slices were cut in a solution containing (in mM): 228 sucrose, 26 NaHCO_3_, 11 glucose, 2.5 KCl, 1 NaH_2_PO_4_, 7 MgSO_4_ and 0.5 CaCl_2_, and recovered in ACSF containing (in mM): 119 NaCl, 26 NaHCO_3_, 11 glucose, 2.5 KCl, 1 NaH_2_PO_4_, 1.3 MgSO_4_ and 2.5 CaCl_2_. Whole-cell recordings were made from presynaptic interneurons using a K-gluconate based internal solution containing (in mM): 143 K-gluconate,10 HEPES, 0.25 EGTA, 2 MgATP, 0.3 Na_3_GTP, 7 phosphocreatine (pH 7.25-7.30; osmolarity 294-298) and from postsynaptic pyramidal cells using an CsCl-based internal solution containing (in mM): 75 CsCl, 68 K-gluconate,10 HEPES, 0.25 EGTA, 2 MgATP, 0.3 Na_3_GTP, 7 phosphocreatine (pH 7.25-7.3; osmolarity 294-298). With this CsCl-based solution, the theoretical reversal potential for Cl^–^ was –14 mV, and IPSCs were inward currents at a holding potential of *–*70 mV. The impedance of patch pipettes was about 3-5 MΩ. Miniature IPSCs (mIPSC) were recorded in pyramidal cells in the presence of 1 μM TTX, 20 μM CNQX and 50 μM D-APV.

For paired recordings (Figure 3F-J and 5G-K), Pv^+^ and SST^+^ interneurons were identified using green fluorescence expressed by AAV-DJ (Ef1**α**-DIO-eYFP). Interneurons in Layer 5 were recorded at current-clamp mode by using K-gluconate based internal solution, and firing pattern was tested by step current injection (-100 to 800 pA). Paired recordings were recorded in Pv^+^ or SST^+^ interneurons (in current-clamp mode) and nearby connected pyramidal (Pyr) cells (in voltage-clamp mode, ~50-100 μm away) by using CsCl based internal solution. Single APs were evoked by injecting current pulses (2 nA, 1 ms). All experiments were performed at 32 °C, using a Multiclamp 700B amplifier (Molecular Devices, LLC). Voltage and current signals were filtered at 10 kHz and sampled at 20 kHz with Digidata 1440 (Molecular Devices, LLC). In PPR experiments in SST cell-Pyr pairs, we utilized optogenetics to simultaneously activate many synaptic inputs from SST^+^ cells by expressing CAG-DIO-ChR2-tdTomato virus. Specifically, CAG-DIO-ChR2-tdtomato (0.5 μl for each hemisphere with a titer of ~5x10^12^) was injected into the mPFC of Nrxn123 cKO/SST-Cre or SST-Cre mice at P21; as a result, ChR2 is only expressed in SST^+^ cells in the mPFC. Pyramidal cell responses to light pulses (0.5 ms, 10-20 mW) were recorded at P35-40. Because ChR2 always evokes complex responses (serrated responses with very different rise time/decay time), the laser intensity was adjusted until a typical monosynaptic response with clear linear rise times and exponential decays was obtained. The amplitude of the first IPSC was usually 100-200 pA, but was dependent on viral expression and light intensity, and cannot be reproducibly quantified. However, paired pulses with different interstimulus intervals could be reliably measured, and the PPR was calculated from such experiments and compared by t-test one by one.

**Two-photon Ca^2+^ imaging**

For Figures 4 and 6, Ca^2+^-imaging was performed with a custom-built laser-scanning system based on an Olympus BX61WI microscope. Two-photon excitation was achieved with a mode-locked Ti:S laser (Chameleon Vision II, Coherent) running at a wavelength of 830 nm (repetition rate: 80 MHz; pulse width: 140 fs). Pv^+^ and SST^+^ interneurons were identified by expression of mCherry from AAV5-Ef1**α**-DIO-mCherry. We recorded Pv^+^ and SST^+^ interneurons with a pipette solution similar to that of paired recordings (K-gluconate based internal solution) but without any additional Ca^2+^-buffer. We utilized Fluo-5F (200 μM) as a Ca^2+^-indicator to monitor Ca^2+^ changes. Alexa Fluor 594 (50 μM) was added into the internal solution for visualization of the cell morphology. The loading of Ca^2+^ indicator took about 20 minutes; Ca^2+^ transients could be then measured. We identified the axons of Pv^+^ and SST^+^ cells by their small diameter, randomly curved collaterals with densely distributed boutons. Axonal terminals were imaged with 5 or 10x digital zoom. APs were elicited by somatic injection of current pulses (1 ms, 2 nA) at 50 Hz. The number of APs varied from 1 to 20. Fluorescence changes were monitored in line scan mode. An individual line scan took 2 ms with 1500 line scans per scan trial. For each bouton and stimulus intensity, the fluorescent signals from 5 trials were averaged to increase the signal/noise ratio. Imaging data were acquired and further analyzed by using MATLAB (MathWorks). Ca^2+^-transients were presented by △G/Gmax calculated from the following equation: △G/G_max_ = [△G/R] /[G_max_/R] (G_max_ is the intensity of 200 μM Fluo-5F in presence of 50 mM Ca^2+^ in ACSF, and R is the intensity of 50 μM Alexa Fluor 594).

**Immunohistochemistry**

Immunohistochemical experiments were performed as previously described (10). Briefly, mice (~P24-P26 or ~P45-55) were anesthetized with isoflurane, perfused with phosphate-buffered saline (PBS) followed by 4% paraformaldehyde (PFA) in 0.1 M PBS via a perfusion pump (2 ml/ min). Perfused brains were postfixed in 4% PFA overnight at 4 °C and then cryoprotected in 30% sucrose (in 1X PBS) for 24 h or 48h at 4 °C. Sagittal cerebellar slices (40 μm) or coronal mPFC slices and hippocampus slices (30 μm) were cryo-sectioned at −20 °C (Leica CM1050). Sections were serially washed with PBS and incubated in blocking solution (0.3% Triton X-100 and 5% goat serum in PBS) for 1 h at room temperature (RT) and incubated overnight at 4 °C with primary antibodies diluted in blocking solution (guinea pig anti-vGluT1, 1:1000; guinea pig anti-vGluT2, 1:500; mouse anti-Calbindin, 1:500; rabbit anti-Calbindin, 1:500; rabbit anti-GFP, 1:1000; rabbit anti-vGAT, 1:500; rabbit anti-Pv, 1:2000). Sections were washed four times (15 min each time) in PBS, treated with secondary antibodies (1:1000, Invitrogen, Alexa 488, Alexa 546, or Alexa 633) at 4 °C overnight or at RT for 1 h, and then washed four additional times (15 min each) again in PBS. Sections were then mounted on superfrost slides and covered with mounting media (Vectashield, Vector Labs).

**Confocal image acquisition and analysis**

Single plane or z-stack images (at 1024 × 1024 resolution) from cerebellar lobules IV/V (areas of sulci excluded) or all layers (L1, L2/3, L5 and L6) of mPFC region or of the hippocampal CA1 region were acquired using a Nikon confocal microscope (A1Rsi) with a 40X (Figure 2E) or 60X oil objective (PlanApo, NA1.4) (all other Figures). All acquisition parameters were kept constant among different conditions within experiments. For each experiment ≥ 8 images per condition were collected and analyzed (mean ± SEM). All unbiased automatic analyses were performed on blinded groups.

In Figures 1 and 2A-C, z-stack images of the cerebellar sections were acquired at 0.5 µm intervals for 30 consecutive sections. Raw ND2 z-stacks were converted to maximum-intensity projections (MIP) in NIS-Elements (Nikon), and all downstream quantification was performed on the projected images. For representative images only, LUT (lookup table in NIS-Elements) adjustments were applied in NIS-Elements for visualization; these adjustments were not used for quantification.

*Background subtraction and segmentation.* For quantitative analysis of synaptic puncta in Figures 3 and S4, the general analysis module was used in NIS-Elements Advanced Research software (Nikon). Prior to thresholding, background fluorescence was measured and reduced using a fixed background-subtraction procedure in NIS-Elements that was applied equally to all images from a given imaging session. Objects were segmented using a binary mask generated by intensity and size thresholding of the relevant channel (e.g., 647 channel for vGAT). Thresholds were defined globally per imaging batch and then applied identically to all images within that batch.  Binary mask settings were optimized and maintained across the images being compared within an experiment. For each ROI, object (or puncta) density was calculated as the number of segmented objects per ROI area, and area fraction was calculated as total segmented binary mask area (in μm²) divided by ROI area.

*Puncta size thresholds.* Puncta size thresholds were set to 0.4–6 µm² for analyses of SynaptoTag and vGluT2⁺ puncta in cerebellar sections (Figure 1 and 2 ). For hippocampal and mPFC analyses, the object size threshold was set to >0.1 μm².  Objects below this size and below intensity threshold were excluded from the binary mask. In Figures 5, 6 and S6, image backgrounds were normalized, and immunoreactive puncta were analyzed with ImageJ software. Puncta size thresholds were set to 0.05–4 μm² for analyses of mPFC and hippocampal sections and 0.1–8 μm² for vGluT2⁺ puncta in cerebellar sections.

*Limitations of threshold-based segmentation.* Synapse density quantifications based on immunostaining are thus subject to limitations both for the area fraction and the synaptic puncta parameters because the results depend entirely on thresholding and reflect multiple parameters (size, antigen content, staining efficacy) that all contribute to the readout. In addition, use of maximum-intensity projections can increase background and can merge puncta/objects that overlap along the z-axis that is poorly resolved, potentially affecting both puncta counts and measured area fraction. Therefore, the reported puncta density and area fraction should be interpreted as relative measures obtained under a standardized segmentation pipeline rather than reflecting absolute numbers. Even as relative measures, effect size vary depending on thresholding (Figure S9).

**STED microscopy**

STED imaging was performed as previously described (11). Immunolabeling was performed using guinea pig anti-vGluT2 (1:500) as the primary antibody, followed by incubation with STAR ORANGE-conjugated secondary antibody (1:500). STED super-resolution images were acquired using a Nikon Ti2-E microscope stand equipped with a STEDYCON confocal and STED module from Abberior Instruments, Inc. Excitation lasers included 488nm (pulsed), 595 nm (pulsed), and 640nm (pulsed). A 775 nm STED depletion laser (pulsed) was used. Detection was performed using time-gated APDs. Images were acquired using a CFI PLAN APO LAMBDA 60X OIL objective with a piezoelectric focusing system. Immersion oil F was used for all imaging. For quantitative analysis, images were acquired using identical settings. Image settings were optimized to ensure that signal was acquired below saturation, with saturation levels indicated using the look-up table. Acquisition laser intensity was scaled ~1.5X higher for STED imaging and the depletion laser was set to enable ~60nm resolution in all channels. Pixel size was automatically determined based on the resolution of the acquired image. For vGluT2 signal analysis, cerebellar sections were immunostained for vGluT2 and imaged for 30-50 consecutive sections. Images were analyzed with Imaris software (Bitplane, Oxford Instruments). The surface rendering module was used to segment vGluT2-positive puncta based on intensity thresholding to minimize non-specific signals. The following parameters were quantified for each region of interest (ROI): (1) volume fraction, defined as the ratio of vGluT2 puncta volume to total tissue volume within the ROI, and (2) mean puncta intensity, measured as the average fluorescence intensity of segmented puncta.

**qRT-PCR and single-cell transcriptional profiling**

For Figure 1B, qRT-PCR was performed as previously described (11).The GFP-positive inferior olive tissue was quickly dissected under a fluorescent stereomicroscope, snap-frozen on dry ice, and then transferred to −80 °C storage until processing. Frozen brain tissues were lysed in Trizol reagent and homogenized using a syringe. Chloroform was then added, and the mixture was vortexed vigorously for 15–20 seconds, followed by incubation on ice for 15 minutes. After centrifugation at 12,000 × g for 15 minutes at 4 °C, the aqueous supernatant was carefully transferred and subjected to RNA extraction using the QIAGEN RNeasy Micro kit. RNA concentration was determined using a NanoDrop 1000 Spectrophotometer and stored at −80 °C until downstream analysis. RNA was converted to cDNA using High Capacity RNA-to-cDNA Kit (ThermoFisher Scientific). Quantitative–PCR was performed with QuantStudio 3 (ThermoFisher Scientific). 10 ng cDNA was used for each reaction, in conjunction with PrimeTime™ Gene Expression Master Mix (IDT ,integrated DNA technologies) and gene specific qRT-PCR probes (IDT ,integrated DNA technologies). The following predesigned assays were used (gene, assay ID): Actb (Mm.PT.39a.22214843.g), Nrxn1 (Mm.PT.58.33547773), Nrxn3 (Mm.PT.58.30432445). Nrxn2 mRNAs were probed using the following assays (gene, primer 1, primer 2, probe): Nrxn2 (5’- CTATACATGGCCTCCCAATGAC-3’, 5’-CGCTGGTGTGTGCTGAA-3’, 5’-FAM/AGTACACGG/ZEN/ATGGACCGC-3’).

For Figure 7, Single-Cell Transcriptional Profiling was performed as previously described (12). Briefly acute brain slices were cut as described above, and patch pipettes were used for cytosol extraction. Samples were then subjected to reverse transcription and target-specific amplification. Pre-amplified cDNAs were then processed for real-time PCR analysis on Biomark 96:96 Dynamic Array according to manufacturer’s protocol (Fluidigm, USA). FAM-dye coupled detection assays were purchased from Integrated DNA Technologies (IDT, USA). To ensure the specificity of the amplification, all assays were tested with dilutions of mouse hippocampal cDNA to verify high efficiency (90-110%) and linear amplification (R^2^>0.96) in every experiment. Expression values were calculated relative to average of ACTB and syt1 assays. Primer and probe sequences used for the single-cell analyses are listed in Table S2. Sequences for the forward primer, internal probe, and reverse primer for assays used in our single-cell analysis. Blank fields are displayed for pre-designed q-PCR assays purchased from IDT.

**Quantifications and statistical analyses**

All data are means ± SEM; numbers of neurons, sections, or ROIs /mice analyzed are shown in the bars or plots. Inter-group comparisons were performed using two-tailed Student’s *t*-test for comparisons of two conditions (bar diagrams) or Kolmogorov-Smirnov test for cumulative distributions, multiple comparisons were analyzed with two-way ANOVA with Bonferroni’s post-test (*p<0.05, **p<0.01, and ***p<0.001). Image backgrounds were normalized using the same settings for all conditions in an experiment, and immunoreactive elements were analyzed with the Nikon analysis software. Test and controls were analyzed on anonymized samples to prevent observer bias. All experiments were performed with at least three independent biological replications.

**Supplementary Figures and Legends**

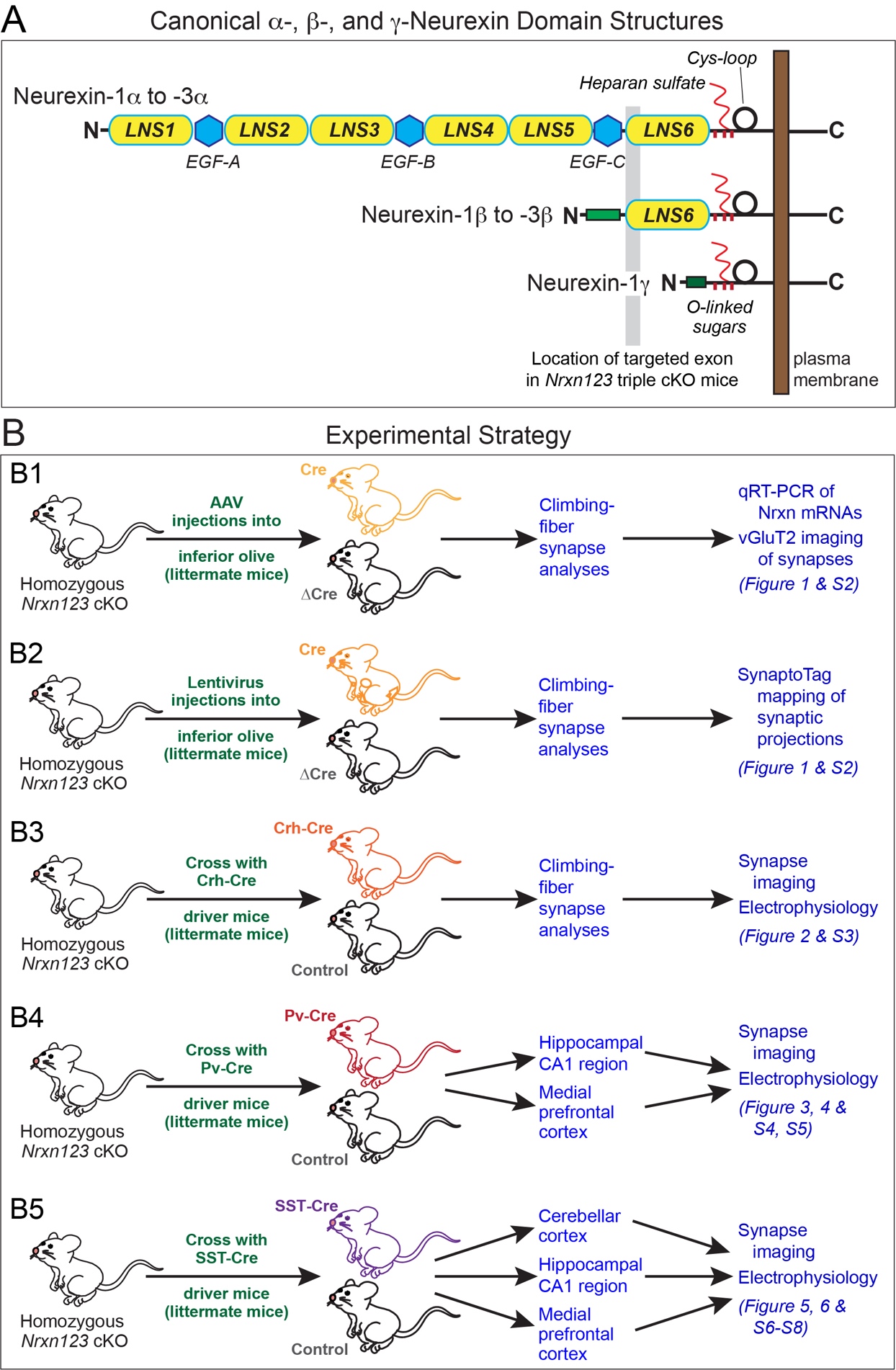

**Figure S1. Domain structures of neurexins and experimental design of the present study**

**A**, Domain structures of neurexins. Three genes (*Nrxn1-3* in mice) encode α- and β-isoforms from independent promoters (13-15), and the Nrxn1 gene additionally encodes a γ-isoform from a third promoter (16). Cre expression in the conditional triple *Nrxn123* KO mice used here (referred to as pan-neurexin deletion) deletes the out-of-frame exon 18 (shaded area) to abolish expression of all neurexin isoforms except for Nrxn1γ. Note that, in addition LNS- and EGF-like domains, neurexins contain a Cys-loop domain followed by a transmembrane region and a cytoplasmic sequence. A subset of neurexins is modified by heparan sulfate as indicated (17), depending on the expression of FAM19A1-4 and CA10/11 proteins that bind to the Cys-loop domain in the secretory pathway and inhibit the heparan sulfate modification (16, 18).

**B**, Schematic of the experimental analyses performed for the current study. We used four different mouse models (B1-B5) to analyze five different synapses in three brain regions (cerebellar climbing-fiber synapses, hippocampal and cortical synapses formed by parvalbumin-positive (Pv^+^) or somatostatin-positive (SST^+^) interneurons). All analyses involved littermate mice whenever possible and were carried out on anonymized samples that blinded the experimenter to the genotype of the sample.

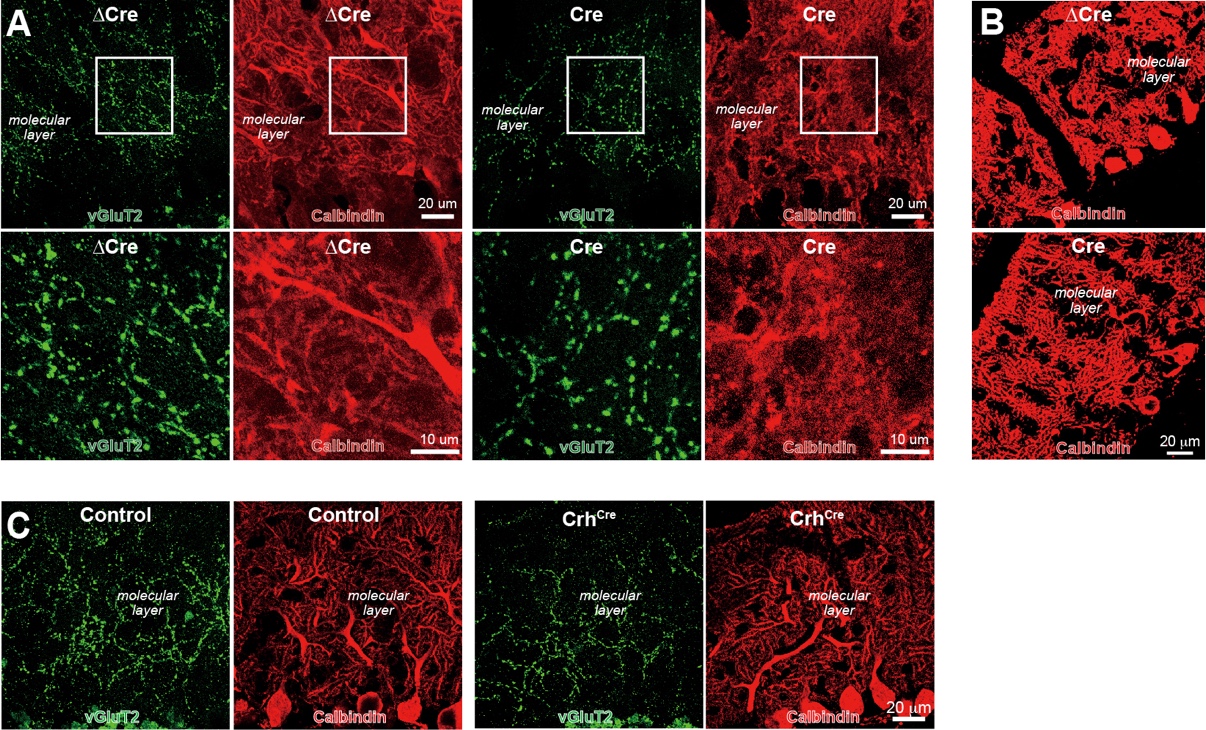

**Figure S2. Representative single-channel images demonstrating that AAV-induced global and lentivirus-induced sparse Cre-mediated pan-neurexin deletions as well as Crh-Cre-induced global pan-neurexin deletions in the inferior olive produce a significant decrease in climbing-fiber synapse density** (complementing Fig. 1, 2)

**A**, Representative red- and green-channel images of the cerebellar cortex of AAV-injected mice visualizing Purkinje cells by calbindin immunofluorescence (red) and climbing-fiber synapses by vGluT2 immunofluorescence (green). The high-magnification images shown below the low-magnification images were taken from the boxed areas. Images are part of the confocal microscopy analysis described in Fig. 1C,D.

**B**, Representative red-channel images of the cerebellar cortex from injected mice visualizing Purkinje cells by calbindin immunofluorescence (red). Images are part of the confocal microscopy analysis described in Fig. 1E, F.

**C**, Representative red- and green-channel images of the cerebellar cortex stained for vGluT2 (green) that is specific for climbing-fiber synapses in the cerebellar cortex and for calbindin (red). Images are part of the confocal microscopy analysis at P28 of the effect of the pan-neurexin deletion in the inferior olive mediated by the Crh-Cre driver line crossed with the triple *Nrxn123* cKO mice (Fig. 2A-C).

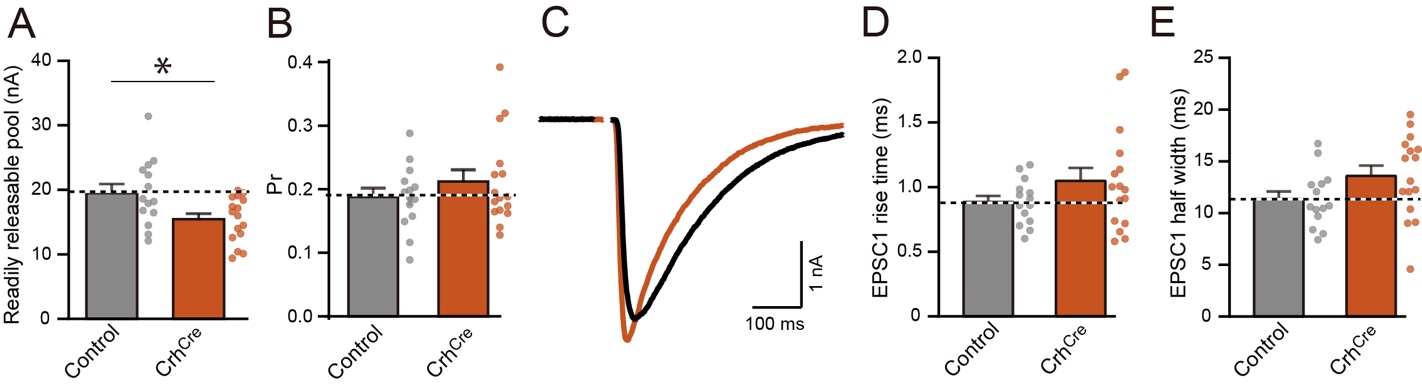

**Figure S3. Further electrophysiological properties of the effect of the presynaptic pan-neurexin deletion in inferior olive neurons on cerebellar climbing-fiber synapses using the Crh-Cre driver mouse line** (complementing Fig. 2)

**A** & **B**, Summary graphs of readily releasable pool size (A) and release probability (Pr) (B) from Purkinje cells in acute cerebellar slices of triple *Nrxn123* cKO mice lacking or harboring a Crh-Cre allele (Control n=14/5, Crh-Cre n=16/5 neurons/mice).

**C**, Representative traces of climbing-fiber EPSC.

**D** & **E**, Summary graphs of EPSC rise time (D) and EPSC half width (E).

Data are mean ± SEM; statistical comparisons were performed with Student’s t-test (*p < 0.05; non-significant comparisons are not labeled).

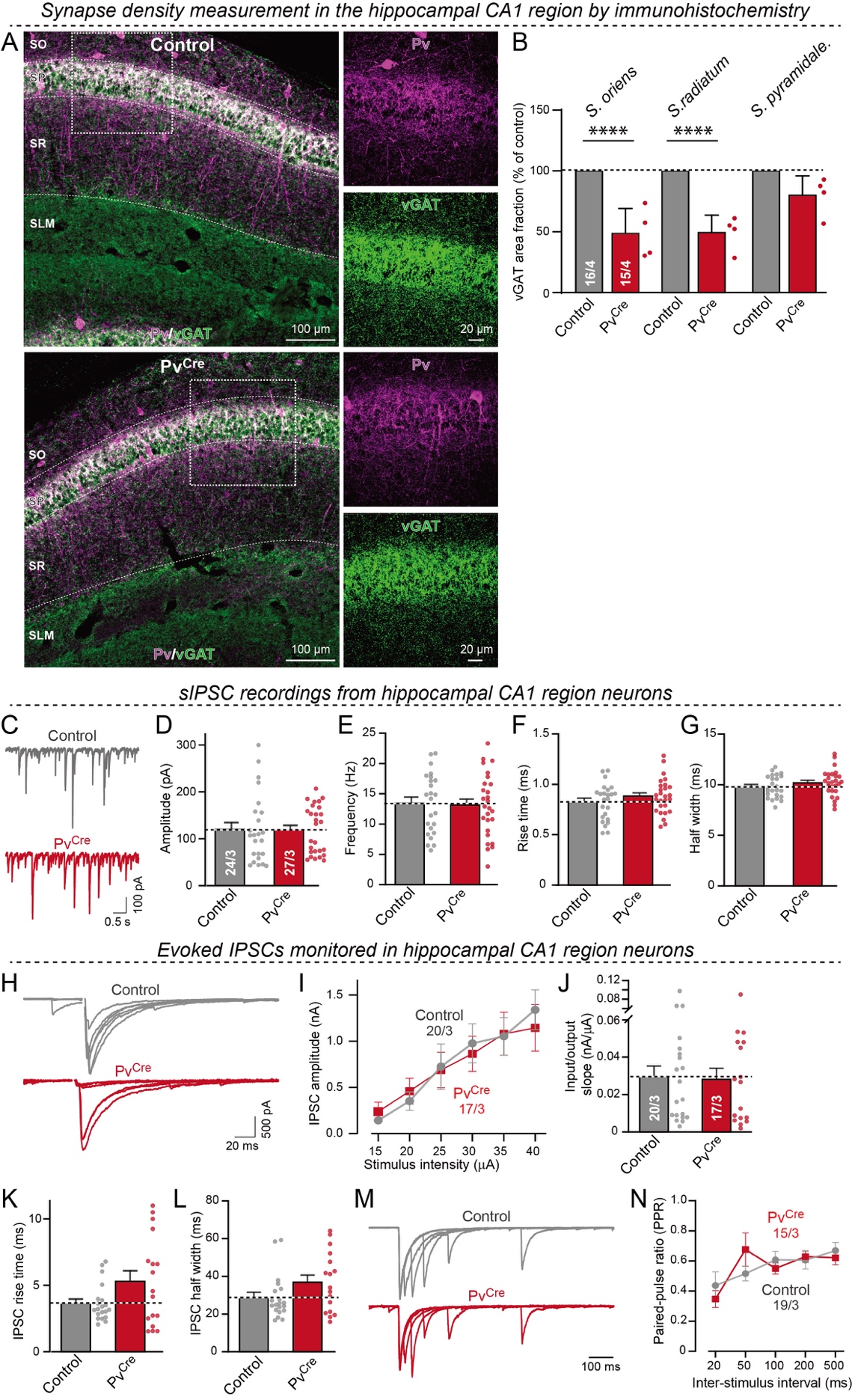

**Figure S4. Pan-neurexin deletions in Pv^+^ interneurons of the hippocampal CA1 region cause an apparent decrease in inhibitory synapse numbers without a detectable inhibitory synapse electrophysiological phenotype**

**A** & **B**, Analysis of the effect of the pan-neurexin deletion in Pv^+^ neurons on the apparent density of inhibitory synapses in the hippocampal CA1 region. Cryostat sections from adult littermate triple *Nrxn123* cKO mice harboring or lacking the Pv-Cre driver allele were analyzed by immunohistochemistry (**A**, representative confocal images of the CA1 region of the hippocampus immunostained for vGAT (green) and parvalbumin (magenta) to visualize inhibitory synapses formed on CA1 pyramidal neurons. Zoomed in confocal single-channel images are represented on the right. **B**, quantifications of the area fraction occupied by inhibitory (vGAT) synapses at CA1 radial layers: SO (Stratum oriens), SR (Stratum radiatum), SP (Stratum pyramidale)).

**C**, Representative spontaneous IPSC (sIPSC) traces recorded from CA1 pyramidal neurons in acute hippocampal slices of triple Nrxn123 cKO mice lacking or harboring a PV-Cre allele.

**D**-**G**, Summary graphs of sIPSC amplitude (D), frequency (E), rise time (F) and half-width (G).

**H**, Representative evoked IPSC traces recorded from CA1 pyramidal neurons in acute hippocampal slices of triple *Nrxn123* cKO mice lacking or harboring a PV-Cre allele.

**I** & **J**, Summary plot of evoked IPSCs as a function of the stimulus intensity and summary graph of the slope of input-output curves of evoked IPSCs.

**K** & **L**, Summary graphs of evoked IPSC rise time (K) and half-width (L).

**M** & **N**, Representative traces of IPSC PPRs recorded from CA1 pyramidal neurons in acute hippocampal slices of triple *Nrxn123* cKO mice lacking or harboring a PV-Cre allele at different inter-stimulus intervals (M) and summary graph of PPRs as a function of the inter-stimulus interval (N).

All numerical data are means ± SEMs (n’s = sections/mice or neurons/mice are indicated in the graphs; statistical evaluations were performed by Student’s t-test (B, D-G, J-L) or two-way ANOVA (I, N) with **** = p<0.0001).

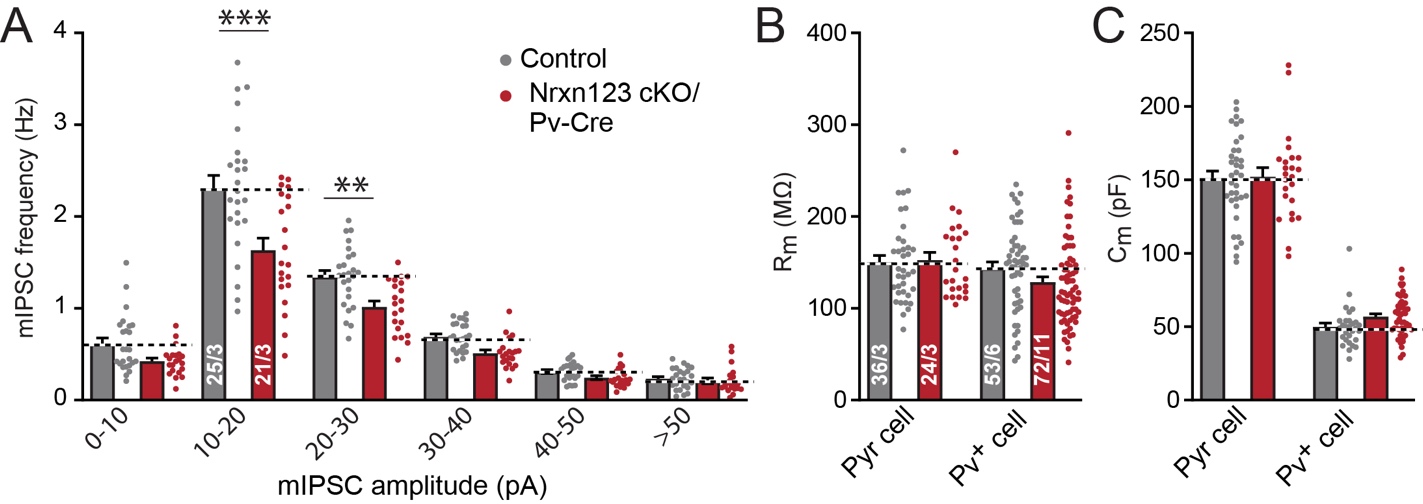

**Figure S5. Additional electrophysiological properties of the effect of the presynaptic pan-neurexin deletion in cortical Pv^+^ neurons** (complementing Fig. 3)

**(A)** Distribution of mIPSC frequencies as a function of mIPSC amplitudes in layer 5 pyramidal neurons from acute mPFC slices of *Nrxn123* cKO and *Nrxn123* cKO/Pv-Cre mice showing that the mIPSC frequency is primarily reduced for events with larger quantal amplitudes by the pan-neurexin deletion. mIPSC events are binned according to the amplitude of individual mIPSC event, and the frequency within each bin was determined by the total number of mIPSC events in the bin divided by the total trial length (300 s). The rationale for this analysis is based on the fact that mIPSCs propagate from synaptic sites that have a wide range of distances from the cell body in which mIPSCs are recorded and may undergo different degrees of attenuation along the way. mIPSCs originating from distal dendrites likely experience greater attenuation and therefore would appear with smaller amplitudes at the soma.

**(B-C)** Summary graph of the membrane resistance (Rm) and capacitance (Cm) of pyramidal (Pyr) and Pv^+^ neurons in *Nrxn123* cKO/Pv-Cre mice vs. controls. Note that pyramidal cells were monitored with a Cs^+^-based internal pipette solution, whereas Pv^+^-interneurons were monitored with a K^+^-based internal solution.

All numerical data are means ± SEMs (n’s = neurons/mice are indicated in the bar graphs); statistical evaluations were performed by Student’s t-test (B, C) or two-way ANOVA followed by Bonferroni *post-hoc* test (A) with * = p<0.05; *** = p<0.001).

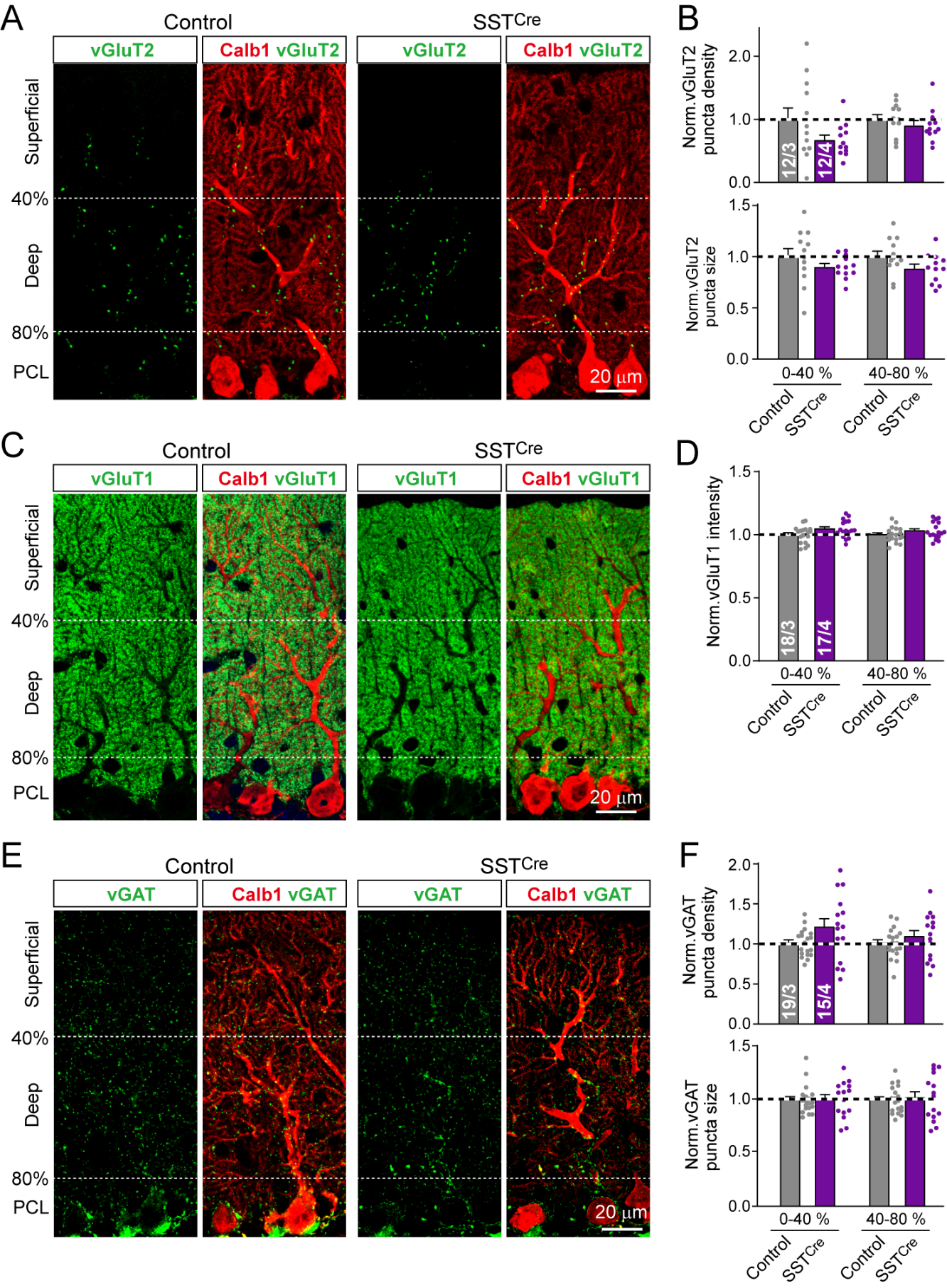

**Figure S6. SST-Cre mediated deletion of all neurexins in SST^+^ interneurons do not change overall synapse number in the cerebellum.**

**(A-B)** Deletion of all neurexins in SST⁺ interneurons do not change the density or size of vGluT2-positive excitatory synaptic puncta corresponding to climbing-fiber synapses in the cerebellar molecular layer (A, representative confocal images of cerebellar cortex sections immunostained for presynaptic vGluT2 and the Purkinje cell marker Calbindin (Calb1) from littermate control *Nrxn123* cKO (left) and *Nrxn123* cKO/SST-Cre mice (right); B, summary graphs of vGluT2-positive puncta density (top) and puncta size (bottom) quantified in the superficial (0–40%) and deep (40–80%) molecular layers of the cerebellar cortex).

(**C-D**) Same as (**A-B**), but for vGluT1-positive excitatory synaptic puncta corresponding to parallel-fiber synapses. Note that only puncta intensity was quantified for analysis.

(**E-F**) Same as (**A-B**), but for vGAT-positive inhibitory synaptic puncta.

All numerical data are means ± SEMs (n’s = sections or slices/mice are indicated in the bar graphs); statistical evaluations were performed by two-way ANOVA followed by Bonferroni *post-hoc* test; non-significant comparisons are not labeled).

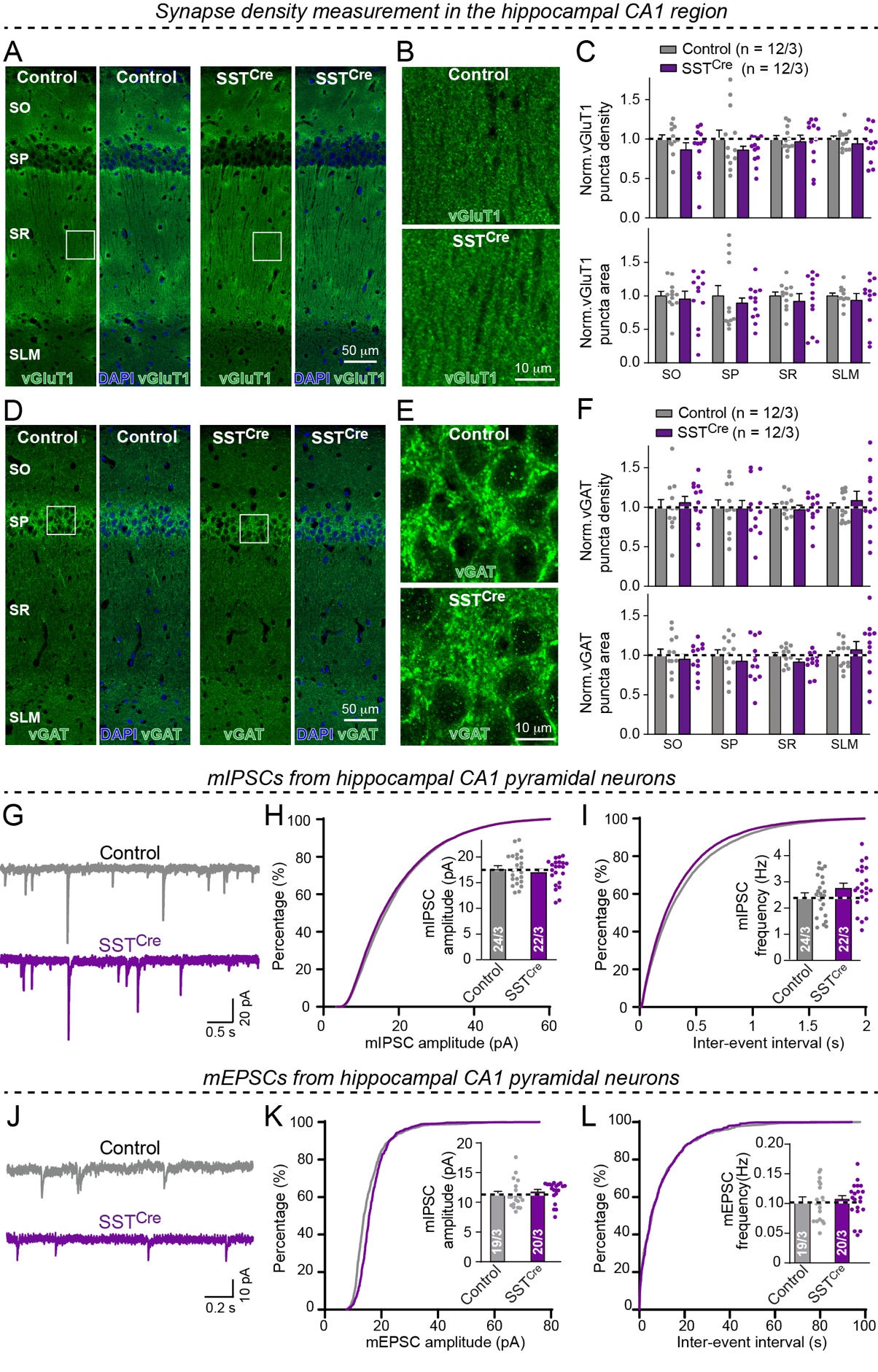

**Figure S7. Pan-neurexin deletions in SST^+^ interneurons do not change overall synapse numbers, mIPSCs, or mEPSCs in the CA1 region of the hippocampus**

**(A-C)** Deletion of all neurexins in SST⁺ interneurons does not change the density and area of vGluT1-positive synaptic puncta in all hippocampal subregions, including the Stratum oriens (SO), Stratum pyramidale (SP), Stratum radiatum (SR), and Stratum lacunosum-moleculare (SLM)(A**,** Representative confocal images of vGluT1 immunostaining showing an overview of the CA1 region from littermate control Nrxn123 cKO (left) and *Nrxn123* cKO/SST-Cre mice (right). B**,** Higher-magnification views of the boxed regions in (A). C**,** Summary graphs of vGluT1⁺ puncta density (top) and area (bottom) quantified in each CA1 subregion).

(**D-F**) Same as (**A-C**), but for vGAT-positive inhibitory synaptic puncta.

(**G-I**) Deletion of all neurexins in SST⁺ interneurons do not change mIPSCs recorded from CA1 pyramidal neurons. (**G**) Representative mIPSC traces. (**H-I**) Cumulative distributions of mIPSC amplitudes (H) and inter-event intervals (I). Insets, summary graphs of mean mIPSC amplitude and frequency.

(**J-L**) Same as (**G-I**), but for mEPSCs.

All numerical data are means ± SEMs (n’s = sections/mice or neurons/mice are indicated in the bar graphs); statistical evaluations were performed by Student’s t-test, two-way ANOVA followed by Bonferroni *post-hoc* test (C, F) or two-tailed Kolmogorov-Smirnov test (cumulative distribution); non-significant comparisons are not labeled. For synapse measurements in the cerebellar cortex, see Figure S3.

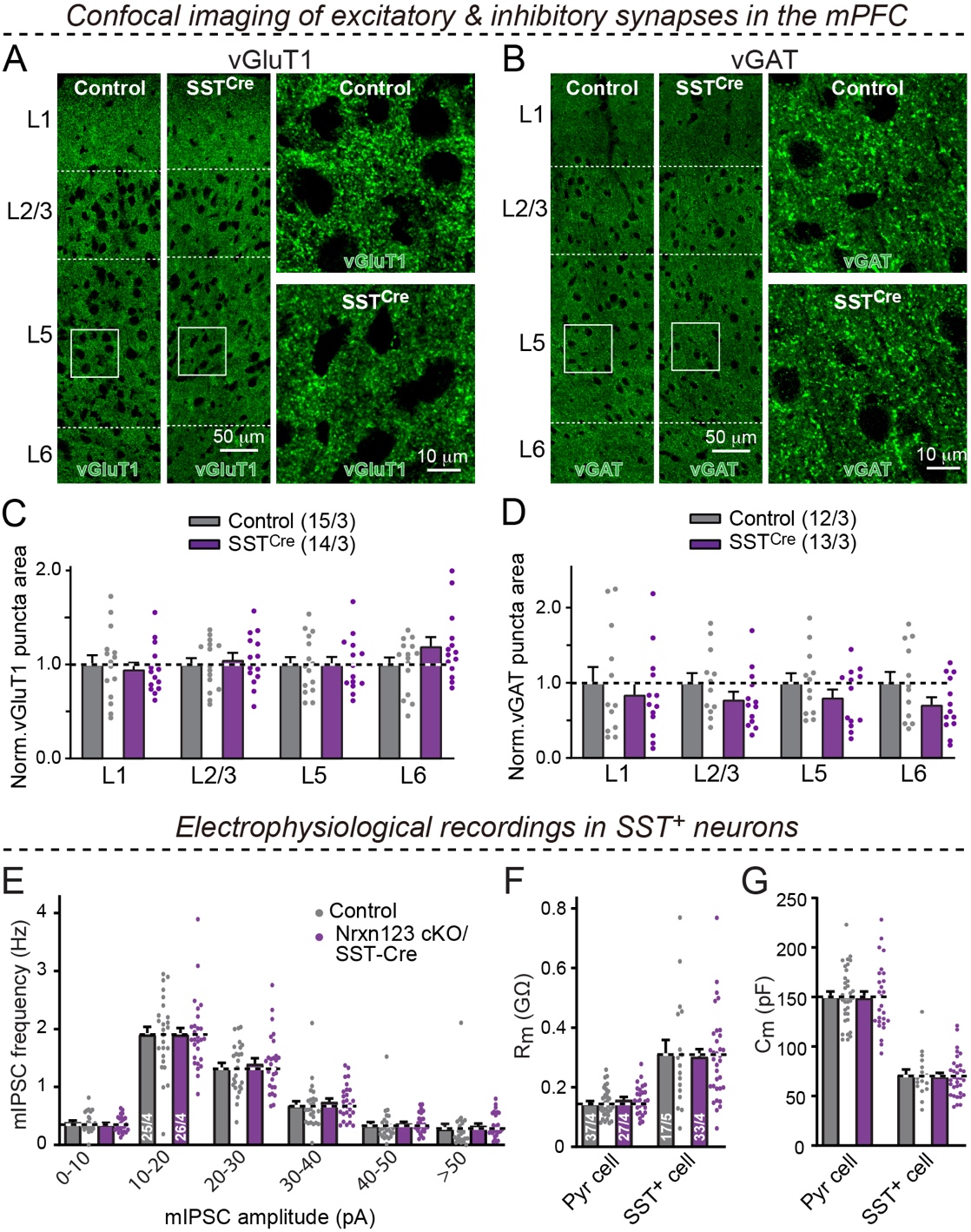

**Figure S8. Representative single-channel images demonstrating that SST-Cre-mediated pan-neurexin deletions do not produce a significant decrease in climbing-fiber synapse density and do** **not change the** **mIPSC frequency recorded in cortical pyramidal neurons as analyzed across quantal amplitudes** (complementing Fig. 5)

**(A** & **B)** Representative green-channel confocal images (L1-L6, cortical layers) from littermate control and SST-Cre mice mPFC sections stained for vGluT1 (A) or vGAT (B). Low-magnification images are shown on the left, high-magnification images taken from the boxed area on the right (corresponding to Fig. 5A, B).

**(C** & **D)** Summary graphs of vGluT1⁺ (C) and vGAT puncta area (D) for each cortical layer, demonstrating that the pan-neurexin deletion does not change the area of vGluT1- and vGAT-positive synaptic puncta in the mPFC (corresponding to Fig. 5C).

**(E)** Distribution of mIPSC frequencies as a function of mIPSC amplitudes in layer 5 pyramidal neurons from acute mPFC slices of *Nrxn123* cKO and *Nrxn123* cKO/SST-Cre mice. All mIPSC events are binned according to the amplitude of individual mIPSC event, and the frequency within each bin was determined by the total number of mIPSC events in that bin divided by the total trial length (300 s).

**(F** & **G)** Summary graph of the membrane resistance (Rm) and capacitance (Cm) of pyramidal (Pyr) cells and SST^+^ cells in *Nrxn123* cKO/SST-Cre vs. SST-Cre controls. Note that pyramidal cells were monitored with a Cs^+^-based internal pipette solution, whereas SST^+^-interneurons were monitored with a K^+^-based internal solution.

All numerical data are means ± SEMs (n’s = sections or slices/mice are indicated in the bar graphs); statistical evaluations were performed by Student’s t-test (B, C) or two-way ANOVA followed by Bonferroni *post-hoc* test (A). Non-significant comparisons are not indicated.

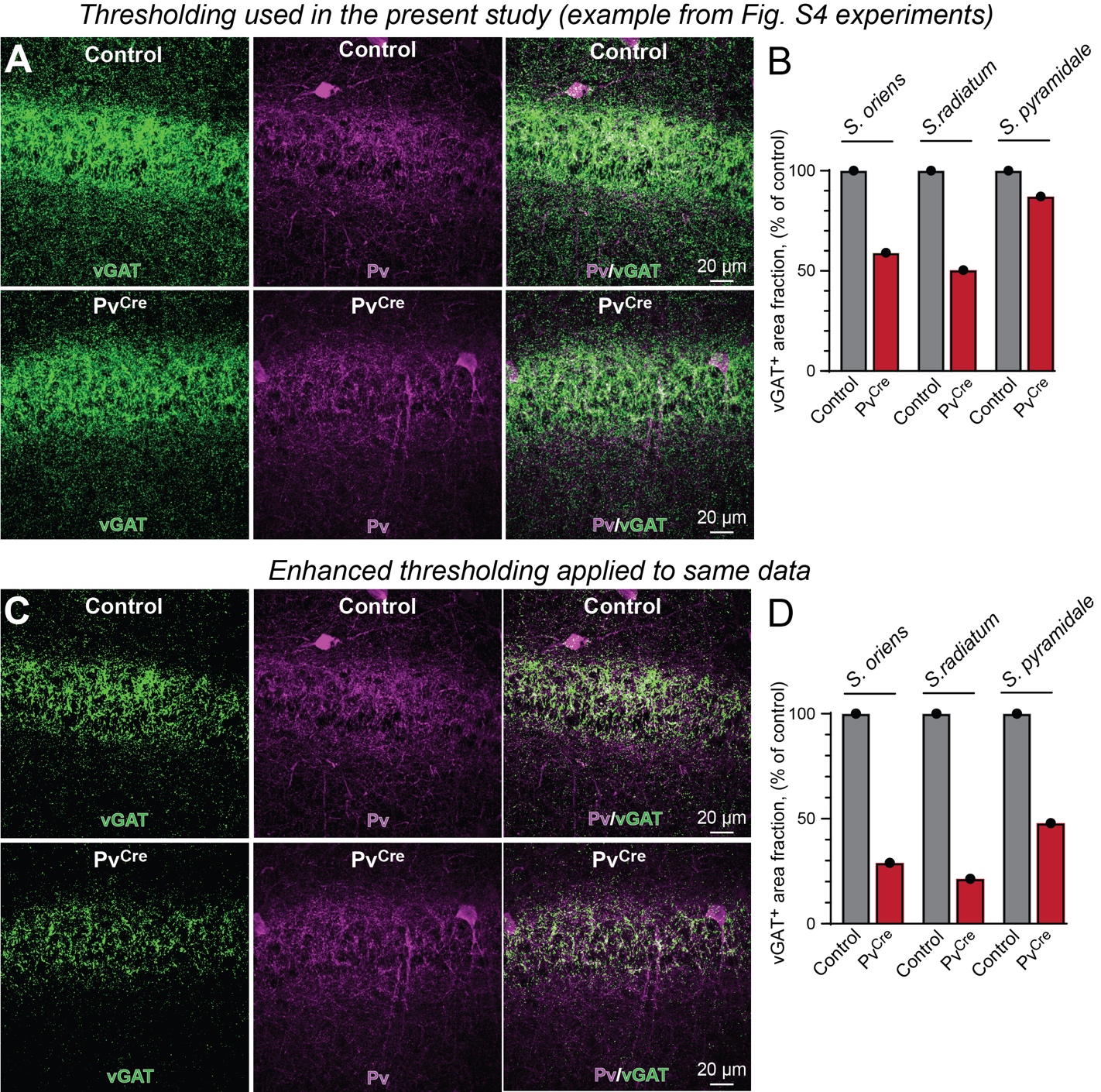

**Figure S9. Illustration of how thresholding affects immunostaining-based synapse-density quantifications** (complementing all morphological synapse studies)

(**A**) Representative maximum-intensity projections images of the CA1 region of the hippocampus from the Figure S4 experiment (Pv, parvalbumin). Images were processed using the same fixed background-subtraction and binary-mask thresholding pipeline as in Figure S4.

(**B**) Quantification of vGAT area fraction (total segmented vGAT mask area normalized to ROI area) across hippocampal strata (str. oriens, str. radiatum, str. pyramidale) using the thresholds shown in (A) for this particular section.

(**C**) The same fields of view as in (A) reprocessed with a higher threshold to allow better vGAT-positive object segmentation, illustrating the sensitivity of segmentation to threshold choice.

(**D**) Quantification of vGAT area fraction from the same dataset using the adjusted thresholds in (C). Across threshold settings, the direction of group differences is preserved, indicating that conclusions are not driven by a specific threshold choice. However, the magnitude differs from (B) as area fraction depends on the number of pixels/objects included in the binary mask, which is sensitive to background fluorescence and the chosen intensity/size thresholds.

Data represent a single experiment to illustrate the effect of thresholding on synapse quantifications. For each sample, the reported total area fraction was calculated as the mean of measurements obtained from three fields of view (FOVs).

**Supplementary Tables**

**Table S1. Key Resources Used in This Study.**

| Reagent or Resource | Source | Identifier |
| --- | --- | --- |
| Alexa Fluor 594 Hydrazide | Invitrogen | Cat# A10438 |
| Fluo-5F, Pentapotassium Salt | Invitrogen | Cat# F14221 |
| Anti-vGluT1, guinea pig | Millipore | AB2301751 |
| Anti-vGluT2, guinea pig | Millipore | AB2665454 |
| Anti-Calbindin, mouse | Sigma | AB476894 |
| Anti-Calbindin, rabbit | Proteintech | 14479-1-AP |
| Anti-GFP, rabbit | Invitrogen | AB221569 |
| Anti-vGAT, rabbit | Synaptic Systems | AB887867 |
| Anti-Pv, rabbit | Swant | AB2315235 |
| Alexa 488, goat anti-mouse | Invitrogen | AB141514 |
| Alexa 488, goat anti-rabbit | Invitrogen | AB221544 |
| Alexa 488, goat anti-guinea pig | Invitrogen | AB142018 |
| Alexa 546, goat anti-mouse | Invitrogen | AB141592 |
| Alexa 546, goat anti-rabbit | Invitrogen | AB143051 |
| Alexa 633, goat anti-mouse | Invitrogen | AB141431 |
| Alexa 633, goat anti-rabbit | Invitrogen | AB2535731 |
| Alexa 633, goat anti-guinea pig | Invitrogen | AB2535757 |
| Vectashield | Vector Labs | AB2336789 |
| MATLAB | MathWorks | SCR001622 |

**Table S2. Primers and probes used for Fluidigm qPCR assays.**

| **Assay Name** | **Gene name** | **Forward Primer** | **Probe** | **Reverse Primer** |
| --- | --- | --- | --- | --- |
| Actin | Actin Beta | AGGTCTTTACGGATGTCAACG | ATTCCATACCCAAGAAGGAAGGCTGG | ATTGGCAACGAGCGGTT |
| Syt1 | Synaptotagmin 1 | ATCATACACAGCCATCACCAG | AGGTGCCATACTCGGAATTAGGTGG | ACCCTCAATCCACTTCTCAATG |
| Sst | Somatostatin | GGCATCATTCTCTGTCTGGTT | AGTTCCTGTTTCCCGGTGGCA | AGACTCCGTCAGTTTCTGC |
| Gad2 | Glutamate decarboxylase 2 | GCCTTGTCTCCTGTGTCATAG | TGCATCAGTCCCTCCTCTCTAACCA | CCTTGCAGTGTTCAGCTCT |
| Pv | Parvalbumin | CTTAGCTTTCAGCCACCAGAG | ATCTTGCCGTCCCCATCCTTGTC | CTGCTAAAGAAACAAAGACGCT |
| Syt2 | Synaptotagmin 2 | Mm.PT.51.12344265 (probe ID) |  |  |
| NRXN1-all | Neurexin 1 | ACTACATCAGTAACTCAGCACAG | CTTCTCCTTGACCACAGCCCCAT | ACAAGTGTCCGTTTCAAATCTTG |
| NRXN1alpha | Neurexin 1 alpha | TCCTCTTAGACATGGGATCAGG | CAACGGGATGGACGGTCAGGTA | GTGTAGGGAGTGCGTAGTG |
| NRXN2alpha | Neurexin 2 alpha | GTCAGCAACAACTTCATGGG | CTTCATCTTCGGGTCCCCTTCCT | AGCCACATCCTCACAGCC |
| NRXN3alpha | Neurexin 3 alpha | GGGAGAACCTGCGAAAGAG | CTGCCGTCATAGCTCAGGATAGATGC | ATGAAGCGGAAGGACACATC |
| CL1/Lphn1 | Latrophilin 1 | GACTGATGCTCTGACTCATGT | TGGGCACACGAAGATGTAAGGGAC | CTGGAACCTACAAATACCTGGA |
| CL2/Lphn2 | Latrophilin 2 | CTCGTGGTGGATATTGTGGTT | TGACCCTGCCCAAGTGCCTAC | TTACGGTATTCCCTGGAGTTTG |
| LRRTM2 | Leucine-rich repeat transmembrane protein 2 | GGCCACTTGAAATGTAAGCC | TGCAGCCTCCAATGTGCTCAGAA | CACTGCGTTGAGTCTGACAA |
| LRRTM3 | Leucine-rich repeat transmembrane protein 3 | CATATGCCAGAAAAGGTTGACAC | AGGCTCCAGGGAATGTGAGATACCT | GAGATGCTGCTGAACGGAA |
| LRRTM4 | Leucine-rich repeat transmembrane protein 4 | GAAAGATAGCACCAGAAACACATC | ACGGAAACCCATCCTTTGTCATCCA | GACCAAATAAGAAGAAAGCTGAAGA |
| NL1 | Neuroligin 1 | GGTTGGGTTTGGTATGGATGA | TGAGGAACTGGTTGATTTGGGTCACC | GATGTTGAGTGCAGTAGTAATGAC |
| NL1-ssA | Neuroligin1 splice site A out | CCAACTGAAGATGATATTCGGG | GGGGTCCCAAACCAGTGATGGTGT | CCAAGACACTCCCATCATACAG |
| Neuroligin2 | Neuroligin 2 | CCGTGTAGAAACAGCATGACC | TCAATCCGCCAGACACCGGATATCCG | TGCCTGTACCTCAACCTCTA |
| Neuroligin3 | Neuroligin 3 | CACTGTCTCGGATGTCTTCA | CCTGTTTCTTAGCGCAGGATCCAT | CCTCTATCTGAATGTGTATGTGC |
| Slitrk2 | SLIT and NTRK-like protein 2 | TCTTTAAGCACCTTCGATGTCA | CCGTCTTCTCCTGATGTGTTTGCCT | GCTAAACTACTGGTGGGACTGAA |
| Slitrk3 | SLIT and NTRK-like protein 3 | GGCTTCATCATTCCTGGTTGA | TGGAGCGAACCTCGGAGATCCT | GACAGAATGACGGGATACAATGC |
| Slitrk4 | SLIT and NTRK-like protein 4 | GTCACTGCCGACGAAGAG | TCCAGAGATAAAGCTGCTCAAAGACCG | CAGATCAAGTAGAGAGCATCGC |
| Slitrk5 | SLIT and NTRK-like protein 5 | GGTCCTGTTCCAAAGTTACTGG | ACGTGCATTTTACCTCTGTTCCCCATCT | TGGGTCGGAGAAAGTTGC |
